## Supplementary Information for "Genetically controlled membrane synthesis in liposomes"

for

by

Duco Blanken,[1] David Foschepoth,[1] Adriana Calaça Serrão, Christophe Danelon*

Department of Bionanoscience, Kavli Institute of Nanoscience, Delft University of Technology, van der Maasweg 9, 2629 HZ, Delft, The Netherlands

[1] Equal contribution

### SUPPLEMENTARY METHODS

#### Cloning of *yfp-spinach* constructs for gene regulation experiments

The *eYFP-LL-spinach* construct was synthesised by Eurogentech (Belgium) and supplied in a pUC57 vector backbone [1]. A lac operator site was introduced via PCR using primers 715 and 716, and *E. coli* TOP10 cells were transformed with the obtained PCR fragment. The T7 promoter in pUC57-T7p-LacO-meYFP-LL-spinach-T7t was substituted with the SP6 promoter using primers 719 and 720 (**Supplementary Table 2**) and the generated pUC57-SP6p-LacO-meYFP-LL-spinach-T7t was used for transformation of TOP10 *E. coli* cells. The identity of the plasmids was confirmed with restriction digestion and Sanger sequencing.

#### Cloning of *fadD* construct

The *fadD* gene was amplified from genomic DNA of *E. coli* K12 with overhangs for Gibson assembly using primers 873 and 874. The vector backbone was amplified from pUC57 with primers 507 and 535. The resulting PCR fragments were purified as described above and used for Gibson assembly. The reaction mixture was subsequently transformed into TOP10 *E. coli* cells. The correct assembly was verified by Sanger sequencing.

#### Production of S30 cell-free lysate

The cell free lysate was prepared according to Hansen et al. [2]. *E. coli* Rosetta2 cells were grown at 37 °C to an $OD_{600}$ of 1.5 in 2 × YTPG broth. After cell growth, all steps were carried out on ice. The cells were collected (3,000 g, 10 min, 4 °C), thoroughly suspended in ice-cold 20% sucrose solution (16 mL for 3 g wet pellet weight) and incubated on ice for 10 min. Cells were then collected (3,000 g, 10 min, 4 °C), resuspended in ice cold Milli-Q (4× wet pellet weight) and immediately spun down (3,000 g, 10 min, 4 °C). Next, cells were again resuspended in ice cold Milli-Q (4× wet pellet weight), allowed to incubate on ice for 10 min and spun down (3,000 g, 10 min, 4 °C). The pellet was then carefully washed twice with ice-cold MilliQ (1.5× volume). The spheroplast pellet was stored at −80 °C. The spheroplasts were thawed and resuspended in ice-cold Milli-Q (0.8× volume). Cells were lysed by 10 cycles of sonication (10 s at 10 µm amplitude followed by 30 s on ice). Cell debris were collected (30,000 g, 30 min, 4 °C) and the supernatant was dialyzed 1× against 50% dialysis buffer (5 mM Tris, 30 mM potassium glutamate, 7 mM magnesium glutamate, 0.5 mM DTT), and 3× 100% dialysis buffer (10 mM Tris, 60 mM potassium glutamate, 14 mM magnesium glutamate, 1 mM DTT).

**Orthogonality assays between SP6 and T7 promoters**

Reaction mixtures consisted of one-third cell lysate from BL21 (DE3) Rosetta2 host strain and two-thirds reaction buffer. The final reaction solution contained 50 mM Hepes (pH 8.0), 2.4 mM guanosine triphosphate, 1 mM each of adenosine triphosphate, cytidine triphosphate and uridine triphosphate, 0.66 mM spermidine, 0.5 mM cyclic adenosine monophosphate, 0.22 mM nicotinamide adenine dinucleotide, 0.17 mM coenzyme A, 20 mM 3-phosphoglyceric acid, 0.045 mM folinic acid, 0.13 mg $mL^{-1}$ transfer ribonucleic acid, 1 mM of each amino acid, 10 mM magnesium glutamate, 66 mM potassium glutamate, 40 U of either T7 or SP6 RNAP (Promega, USA), 1 mM DFHBI [3] and cell lysate, contributing an additional 5 mM magnesium glutamate and 20 mM potassium glutamate. 7 nM of plasmid DNA (either pUC57-T7p-LacO-meYFP-LL-spinach-T7t or pUC57-SP6p-LacO-meYFP-LL-spinach-T7t was added last to trigger transcription/translation.

### SUPPLEMENTARY NOTES

#### Supplementary Note 1: Rational for gene orientation in pGEMM7

In a previous version of the plasmid (pGEMM6) both *pgsA* and *pgpA* genes were under control of an SP6 promoter but orientated in the same direction as the other genes on the plasmid (**Supplementary Fig. 4a**). We investigated whether PG production was conditional to the presence of SP6 RNAP. Plasmid pGEMM6 was expressed in PURE system containing LUVs and lipid synthesis was assessed by mass spectrometry. DOPG was produced also in the absence of SP6 RNAP (**Supplementary Fig. 4b**), showing the lack of transcriptional orthogonality. We suspected that read-through transcription might be the cause of unintended production of *pgsA* and *pgpA* transcripts, eventually leading to PG synthesis. To remedy this problem, we designed pGEMM7, where the two genes under SP6 promoter control were flipped in the opposite reading direction (**Supplementary Fig. 1**). As expected, expression of pGEMM7 resulted in no pgsA protein and no PG phospholipid production when the SP6 RNAP was omitted (main text **Fig. 1b,c**).

#### Supplementary Note 2: Absence of PgpA specific signal in MS data

We were not able to detect a suitable peptide for PgpA. Therefore, PgsA was used as a reporter of transcriptional activation of the PgsA-PgpA branch upon SP6 RNAP addition (main text **Fig. 1a,b**). As an alternative to PgpA, PgpC (Uniprot: P0AD42) can catalyze the same enzymatic reaction as PgpA (**Supplementary Fig. 8**). Four peptides of PgpC were detected with our protocol for trypsin digestion and LC-MS/MS as shown in **Supplementary Fig. 7**. They are listed in **Supplementary Table 5**.

#### Supplementary Note 3: Kinetic analysis of LactC2-eGFP association to PS-producing liposomes

Phenomenological fitting was applied to extract kinetic parameters from the time traces reported in main text **Fig. 5g** (**Supplementary Fig. 19**). The following sigmoid equation was used:

$$y = k' + k\,\frac{t^n}{t^n + K^n} \qquad \text{(Eq. 1)}$$

Where $t$ is the time in minutes, and $y$ is the LactC2-eGFP fluorescence signal at the rim of a liposome at a given time point. $k$, $k'$, $K$, and $n$ are fitting parameters. This equation has previously been used to fit fluorescence measurements of cell-free gene expression of YFP in liposomes [4], and mass spectrometry measurements of cell-free Min protein expression [5]. The plateau time, i.e. the time until LactC2-eGFP fluorescence stops increasing, was defined as:

$$T_{plateau} = \frac{2K}{n} + K. \qquad \text{(Eq. 2)}$$

The rate, which is the steepness at time $t$ = $K$, was defined as:

$$\text{rate} = \frac{kn}{4K}. \quad \text{(Eq. 3)}$$

This apparent rate captures the rate of transcription and translation, the enzyme kinetics leading to the synthesis of PS, the membrane incorporation of PS, as well as the binding kinetics of LactC2-eGFP to PS, as illustrated in **Supplementary Fig. 19a**.

### SUPPLEMENTARY TABLES

**Supplementary Table 1: List of plasmids used in this study**

| Plasmid | Size (kb) | Encoded genes |
|---|---|---|
| pGEMM7 | 10.5 | *plsB*, *plsC*, *cdsA*, *pssA*, *psd*, *pgsA*, *pgpA* |
| pGEMM6 | 10.5 | *plsB*, *plsC*, *cdsA*, *pssA*, *psd*, *pgsA*, *pgpA* |
| pUC57-T7p-LacO-meYFP-LL-spinach-T7t | 3.8 | *meYFP* |
| pUC57-SP6p-LacO-meYFP-LL-spinach-T7t | 3.8 | *meYFP* |
| pUC57-T7p-fadD | 4.5 | *fadD* |

### Supplementary Table 2: List of primers used in this study

| Primer | Sequence, 5' to 3' | Linker site | Target |
|---|---|---|---|
| 25 ChD | GATGCTGTAGGCATAGGCTTGG | | |
| 91 ChD | AAAAAACCCCTCAAGACCCGTTTAGAGG | | |
| 288 ChD | CGATGCGTCCGGC | | |
| 310 ChD | GGATCTCGACGCTCTCCCTTATG | | |
| 397 ChD | CCTCTAGAAATAATTTTGTTTAACTTTAAGAAGG | | |
| 471 ChD | CAAAGCCCGAAAGGAAGCTGA | | |
| 507 ChD | ATGTATATCTCCTTCTTAAAGTTAAACAAAATTATTTCTAGAGG | | pUC57 |
| 535 ChD | TAGCATAACCCCTTGGGGCCTCTAAAC | | pUC57 |
| 628 ChD | TGGGCCCGTAACAAAATCCTCCCAATAAGCTAATACGACTCACTATAGG | 1 top | *plsB* |
| 629 ChD | ACCAGGGCTGTTCAACCGACGCTCACGGGGCAAAAAACCCCTCAAGACC | 2 bottom | *plsB* |
| 630 ChD | CCCCGTGAGCGTCGGTTGAACAGCCCTGGTTAATACGACTCACTATAGG | 2 top | *plsC* |
| 631 ChD | AGCGATATATTCGGGCTTCTGGTCGGGCCGCAAAAAACCCCTCAAGACC | 3 bottom | *plsC* |
| 632 ChD | CGGCCCGACCAGAAGCCCGAATATATCGCTTAATACGACTCACTATAGG | 3 top | *cdsA* |
| 633 ChD | AGGTAACGCACCCCGGCCCAAGAGCCGTAACAAAAAACCCCTCAAGACC | 4 bottom | *cdsA* |
| 634 ChD | TTACGGCTCTTGGGCCGGGGTGCGTTACCTTAATACGACTCACTATAGG | 4 top | *pssA* |
| 635 ChD | AGGAATTAACGGACGGCCTCGATTTCTGCACAAAAAACCCCTCAAGACC | 5 bottom | *pssA* |
| 636 ChD | TGCAGAAATCGAGGCCGTCCGTTAATTCCTTAATACGACTCACTATAGG | 5 top | *psd* |
| 715 ChD | GGAATTGTGAGCGGATAACAATTCCCCTCTAGAAATAATTTTGTTTAACTTT | | |
| 716 ChD | GAATTGTTATCCGCTCACAATTCCGGTCTCCCTATAGTGAGTCG | | |
| 719 ChD | ATTTAGGTGACACTATAGAAGAACCGGAATTGTGAGCGGA | | |
| 720 ChD | TCTTCTATAGTGTCACCTAAATATTTCGCATCTAGATGCATTCGC | | |
| 723 ChD | TCGATATTGGGGACTTCTCAAATCTCGTCACAAAAAACCCCTCAAGACC | SHR18 bottom | *psd* |
| 848 ChD | TATACATATGGGCAGCAGCCATCATCATCATCATCACAGCAGCGGCCTGGTGCCGCGCG GCAGCCATATGGTGAGCAAGGGCGAGG | | |
| 849 ChD | AACTCAGCTTCCTTTCGGGCTTTGCTAACAGCCCAGCAGCTCC | | |
| 850 ChD | GATGATGGCTGCTGCCCATATGTATATCTCCTTCTTAAAGTTAAAC | | |
| 851 ChD | CCTGCAGGCCTTAGGCTCGAGCGGCCGCTGAGGACTAGTATTTAGGTGACACTATAGA | 13 top | *pgpA* |
| 852 ChD | TGACGAGATTTGAGAAGTCCCCAATATCGACAAAAAACCCCTCAAGACC | SHR18 top | *pgsA* |
| 829 ChD | TGACGAGATTTGAGAAGTCCCCAATATCGAATTTAGGTGACACTATAGAAGAACCCCTCT AGAAATAATTTTGTTTAACTTTAAGAAGG | 13 bottom | pUC19 |
| 830 ChD | GCTTATTGGGAGGATTTTGTTACGGGCCCAGAATCAGGGGATAACGCAGG | 1 bottom | pUC19 |
| 818 ChD | GTGTCAGTCAATCACGGGCGGGTCCACTACCAAAAAACCCCTCAAGACCCGTTTAG | 30 bottom | *pgpA* |

| | | | |
|---|---|---|---|
| 819 ChD | GTAGTGGACCCGCCCGTGATTGACTGACACATTTAGGTGACACTATAGAAGAATCCCCT CTAGAAATAATTTTGTTTAAC | 30 top | *pgsA* |
| 873 ChD | GTTTAACTTTAAGAAGGAGATATACATATGAAGAAGGTTTGGCTTAACCGTTATCC | | *fadD* |
| 874 ChD | CCCGTTTAGAGGCCCCAAGGGGTTATGCTAGTCAGGCTTTATTGTCCACTTTGCCG | | *fadD* |

**Supplementary Table 3: Pipetting scheme of Gibson assembly to obtain pGEMM7**

| Part | Size (kb) | Ratio length to backbone | Mass DNA added into reaction (ng) |
|---|---|---|---|
| pUC19 | 2 | 1 | 100 |
| *pgpA* | 0.8 | 0.4 | 40 |
| *pgsA* | 0.8 | 0.4 | 40 |
| *plsB* | 2.6 | 1.3 | 130 |
| *plsC* | 0.9 | 0.45 | 45 |
| *cdsA* | 1.2 | 0.6 | 60 |
| *pssA* | 1.4 | 0.7 | 70 |
| *psd* | 1 | 0.5 | 50 |

**Supplementary Table 4: Full list of the tryptic peptides detected by LC-MS/MS in this study.** The first column shows the uniprot accession number as well as the gene name. The peptide sequence is shown in column 2 followed by the standard type used (internal for absolute quantification, global for normalization). The start and end positions of the peptide in the wild-type gene are indicated. In the last two columns, the predicted and measured retention time values are reported.

| Protein (Uniprot No.) | Peptide | Standard type | Begin #AA | End #AA | Predicted retention time (min) | Average measured retention time (min) |
|---|---|---|---|---|---|---|
| PlsB (P0A7A7) | LLNLPLSILVK | | 10 | 20 | 17.87 | 20.31 |
| PlsB (P0A7A7) | FSPSVSLR | | 166 | 173 | 7.77 | 8.09 |
| PlsB (P0A7A7) | LAAVGPR | | 202 | 208 | 2.58 | 2.9 |
| PlsB (P0A7A7) | DTIGDIIILPR | Internal standard | 589 | 599 | 16.94 | 16.81 |
| PlsC (P26647) | LAPLFGLK | | 44 | 51 | 13.66 | 13.48 |
| PlsC (P26647) | GLLPFK | | 154 | 159 | 10.14 | 10.49 |
| CdsA (P0ABG1) | DSGHLIPGHGGILDR | | 248 | 262 | 10.4 | 8.1 |
| PssA (P23830) | GILNALYEAK | | 65 | 74 | 11.93 | 12.49 |
| PssA (P23830) | DLQSIADYPVK | Internal standard | 419 | 429 | 9.57 | 11 |
| Psd (P0A8K1) | LSLQYILPK | | 6 | 14 | 13.64 | 14.89 |
| Psd (P0A8K1) | LVIDLFVK | | 35 | 42 | 16.2 | 15.41 |
| PgsA (P0ABF8) | EIIISALR | | 101 | 108 | 12.73 | 11.36 |
| PgsA (P0ABF8) | SSVAVSWIGK | Internal standard | 118 | 127 | 11.03 | 10.34 |
| EF-Tu (P0CE47) | TTLTAAITTVLAK | Global standard | 25 | 37 | 13.94 | 16.65 |
| EF-Tu (P0CE47) | GITINTSHVEYDTPTR | | 59 | 74 | 9.11 | 9.56 |

### Supplementary Table 5: Transitions of the MS/MS measurements for the proteomic analysis

| Protein (Uniprot No.) | Peptide | Precursor ion *m/z* | MS1 Res | Product ion *m/z* | MS2 Res | Dwell time (ms) | Fragmentor (V) | Collision energy (V) | Ion |
|---|---|---|---|---|---|---|---|---|---|
| PlsB (P0A7A7) | LLNLPLSILVK | 611.9103 | Unit | 769.5182 | Unit | 20 | 130 | 20 | y7 |
| PlsB (P0A7A7) | LLNLPLSILVK | 611.9103 | Unit | 227.1754 | Unit | 20 | 130 | 20 | b2 |
| PlsB (P0A7A7) | LLNLPLSILVK | 611.9103 | Unit | 341.2183 | Unit | 20 | 130 | 20 | b3 |
| PlsB (P0A7A7) | LLNLPLSILVK | 611.9103 | Unit | 454.3024 | Unit | 20 | 130 | 20 | b4 |
| PlsB (P0A7A7) | FSPSVSLR | 446.748 | Unit | 658.3883 | Unit | 20 | 130 | 14.8 | y6 |
| PlsB (P0A7A7) | FSPSVSLR | 446.748 | Unit | 375.235 | Unit | 20 | 130 | 14.8 | y3 |
| PlsB (P0A7A7) | FSPSVSLR | 446.748 | Unit | 288.203 | Unit | 20 | 130 | 14.8 | y2 |
| PlsB (P0A7A7) | FSPSVSLR | 446.748 | Unit | 235.1077 | Unit | 20 | 130 | 14.8 | b2 |
| PlsB (P0A7A7) | LAAVGPR | 342.2136 | Unit | 499.2987 | Unit | 20 | 130 | 11.6 | y5 |
| PlsB (P0A7A7) | LAAVGPR | 342.2136 | Unit | 428.2616 | Unit | 20 | 130 | 11.6 | y4 |
| PlsB (P0A7A7) | LAAVGPR | 342.2136 | Unit | 329.1932 | Unit | 20 | 130 | 11.6 | y3 |
| PlsB (P0A7A7) | LAAVGPR | 342.2136 | Unit | 185.1285 | Unit | 20 | 130 | 11.6 | b2 |
| PlsB (P0A7A7) | DTIGDIIILPR | 613.3612 | Unit | 896.5564 | Unit | 20 | 130 | 20 | y8 |
| PlsB (P0A7A7) | DTIGDIIILPR | 613.3612 | Unit | 611.4239 | Unit | 20 | 130 | 20 | y5 |
| PlsB (P0A7A7) | DTIGDIIILPR | 613.3612 | Unit | 498.3398 | Unit | 20 | 130 | 20 | y4 |
| PlsB (P0A7A7) | DTIGDIIILPR | 613.3612 | Unit | 385.2558 | Unit | 20 | 130 | 20 | y3 |
| PlsB (P0A7A7) | DTIGDIIILPR | 613.3612 | Unit | 272.1717 | Unit | 20 | 130 | 20 | y2 |
| PlsB (P0A7A7) | DTIGDIIILPR.heavy | 618.3653 | Unit | 906.5646 | Unit | 20 | 130 | 20 | y8 |
| PlsB (P0A7A7) | DTIGDIIILPR.heavy | 618.3653 | Unit | 621.4322 | Unit | 20 | 130 | 20 | y5 |
| PlsB (P0A7A7) | DTIGDIIILPR.heavy | 618.3653 | Unit | 508.3481 | Unit | 20 | 130 | 20 | y4 |
| PlsB (P0A7A7) | DTIGDIIILPR.heavy | 618.3653 | Unit | 395.264 | Unit | 20 | 130 | 20 | y3 |
| PlsB (P0A7A7) | DTIGDIIILPR.heavy | 618.3653 | Unit | 282.18 | Unit | 20 | 130 | 20 | y2 |
| PlsC (P26647) | LAPLFGLK | 429.776 | Unit | 674.4236 | Unit | 20 | 130 | 14.3 | y6 |
| PlsC (P26647) | LAPLFGLK | 429.776 | Unit | 464.2867 | Unit | 20 | 130 | 14.3 | y4 |
| PlsC (P26647) | LAPLFGLK | 429.776 | Unit | 337.7154 | Unit | 20 | 130 | 14.3 | y6 |
| PlsC (P26647) | LAPLFGLK | 429.776 | Unit | 185.1285 | Unit | 20 | 130 | 14.3 | b2 |
| PlsC (P26647) | GLLPFK | 337.7154 | Unit | 504.318 | Unit | 20 | 130 | 11.5 | y4 |
| PlsC (P26647) | GLLPFK | 337.7154 | Unit | 391.234 | Unit | 20 | 130 | 11.5 | y3 |

| | | | | | | | | | |
|---|---|---|---|---|---|---|---|---|---|
| PlsC (P26647) | GLLPFK | 337.7154 | Unit | 171.1128 | Unit | 20 | 130 | 11.5 | b2 |
| PlsC (P26647) | GLLPFK | 337.7154 | Unit | 284.1969 | Unit | 20 | 130 | 11.5 | b3 |
| CdsA (P0ABG1) | DSGHLIPGHGGILDR | 515.2707 | Unit | 630.3570 | Unit | 20 | 130 | 8.15 | Y6 |
| CdsA (P0ABG1) | DSGHLIPGHGGILDR | 515.2707 | Unit | 397.1466 | Unit | 20 | 130 | 8.15 | b4 |
| CdsA (P0ABG1) | DSGHLIPGHGGILDR | 515.2707 | Unit | 510.2306 | Unit | 20 | 130 | 8.15 | b5 |
| CdsA (P0ABG1) | DSGHLIPGHGGILDR | 515.2707 | Unit | 623.3147 | Unit | 20 | 130 | 8.15 | b6 |
| PssA (P23830) | GILNALYEAK | 546.3084 | Unit | 921.504 | Unit | 20 | 130 | 17.9 | y8 |
| PssA (P23830) | GILNALYEAK | 546.3084 | Unit | 808.4199 | Unit | 20 | 130 | 17.9 | y7 |
| PssA (P23830) | GILNALYEAK | 546.3084 | Unit | 623.3399 | Unit | 20 | 130 | 17.9 | y5 |
| PssA (P23830) | DLQSIADYPVK | 624.8272 | Unit | 892.4775 | Unit | 20 | 130 | 20.4 | y8 |
| PssA (P23830) | DLQSIADYPVK | 624.8272 | Unit | 692.3614 | Unit | 20 | 130 | 20.4 | y6 |
| PssA (P23830) | DLQSIADYPVK | 624.8272 | Unit | 506.2973 | Unit | 20 | 130 | 20.4 | y4 |
| PssA (P23830) | DLQSIADYPVK | 624.8272 | Unit | 357.1769 | Unit | 20 | 130 | 20.4 | b3 |
| PssA (P23830) | DLQSIADYPVK.heavy | 628.8343 | Unit | 900.4917 | Unit | 20 | 130 | 20.4 | y8 |
| PssA (P23830) | DLQSIADYPVK.heavy | 628.8343 | Unit | 700.3756 | Unit | 20 | 130 | 20.4 | y6 |
| PssA (P23830) | DLQSIADYPVK.heavy | 628.8343 | Unit | 514.3115 | Unit | 20 | 130 | 20.4 | y4 |
| PssA (P23830) | DLQSIADYPVK.heavy | 628.8343 | Unit | 357.1769 | Unit | 20 | 130 | 20.4 | b3 |
| Psd (P0A8K1) | LSLQYILPK | 537.8315 | Unit | 633.397 | Unit | 20 | 130 | 17.7 | y5 |
| Psd (P0A8K1) | LSLQYILPK | 537.8315 | Unit | 357.2496 | Unit | 20 | 130 | 17.7 | y3 |
| Psd (P0A8K1) | LSLQYILPK | 537.8315 | Unit | 244.1656 | Unit | 20 | 130 | 17.7 | y2 |
| Psd (P0A8K1) | LSLQYILPK | 537.8315 | Unit | 831.4975 | Unit | 20 | 130 | 17.7 | b7 |
| Psd (P0A8K1) | LVIDLFVK | 473.8022 | Unit | 833.5131 | Unit | 20 | 130 | 15.7 | y7 |
| Psd (P0A8K1) | LVIDLFVK | 473.8022 | Unit | 734.4447 | Unit | 20 | 130 | 15.7 | y6 |
| Psd (P0A8K1) | LVIDLFVK | 473.8022 | Unit | 506.3337 | Unit | 20 | 130 | 15.7 | y4 |
| Psd (P0A8K1) | LVIDLFVK | 473.8022 | Unit | 213.1598 | Unit | 20 | 130 | 15.7 | b2 |
| PgsA (P0ABF8) | EIIISALR | 457.7871 | Unit | 559.3562 | Unit | 20 | 130 | 15.2 | y5 |
| PgsA (P0ABF8) | EIIISALR | 457.7871 | Unit | 446.2722 | Unit | 20 | 130 | 15.2 | y4 |
| PgsA (P0ABF8) | EIIISALR | 457.7871 | Unit | 359.2401 | Unit | 20 | 130 | 15.2 | y3 |
| PgsA (P0ABF8) | EIIISALR | 457.7871 | Unit | 356.218 | Unit | 20 | 130 | 15.2 | b3 |
| PgsA (P0ABF8) | SSVAVSWIGK | 517.2875 | Unit | 760.4352 | Unit | 20 | 130 | 17 | y7 |
| PgsA (P0ABF8) | SSVAVSWIGK | 517.2875 | Unit | 689.3981 | Unit | 20 | 130 | 17 | y6 |

| | | | | | | | | | |
|---|---|---|---|---|---|---|---|---|---|
| PgsA (P0ABF8) | SSVAVSWIGK | 517.2875 | Unit | 590.3297 | Unit | 20 | 130 | 17 | y5 |
| PgsA (P0ABF8) | SSVAVSWIGK | 517.2875 | Unit | 345.1769 | Unit | 20 | 130 | 17 | b4 |
| PgsA (P0ABF8) | SSVAVSWIGK.heavy | 521.2946 | Unit | 768.4494 | Unit | 20 | 130 | 17 | y7 |
| PgsA (P0ABF8) | SSVAVSWIGK.heavy | 521.2946 | Unit | 697.4123 | Unit | 20 | 130 | 17 | y6 |
| PgsA (P0ABF8) | SSVAVSWIGK.heavy | 521.2946 | Unit | 598.3439 | Unit | 20 | 130 | 17 | y5 |
| PgsA (P0ABF8) | SSVAVSWIGK.heavy | 521.2946 | Unit | 345.1769 | Unit | 20 | 130 | 17 | b4 |
| EF-TU (P0CE47) | TTLTAAITTVLAK | 652.3952 | Unit | 887.556 | Unit | 20 | 130 | 21.2 | y9 |
| EF-TU (P0CE47) | TTLTAAITTVLAK | 652.3952 | Unit | 816.5189 | Unit | 20 | 130 | 21.2 | y8 |
| EF-TU (P0CE47) | TTLTAAITTVLAK | 652.3952 | Unit | 745.4818 | Unit | 20 | 130 | 21.2 | y7 |
| EF-TU (P0CE47) | TTLTAAITTVLAK | 652.3952 | Unit | 632.3978 | Unit | 20 | 130 | 21.2 | y6 |
| EF-TU (P0CE47) | GITINTSHVEYDTPTR | 601.9672 | Unit | 752.3573 | Unit | 20 | 130 | 16.9 | y6 |
| EF-TU (P0CE47) | GITINTSHVEYDTPTR | 601.9672 | Unit | 474.2671 | Unit | 20 | 130 | 16.9 | y4 |
| EF-TU (P0CE47) | GITINTSHVEYDTPTR | 601.9672 | Unit | 710.3286 | Unit | 20 | 130 | 16.9 | y12 |
| EF-TU (P0CE47) | GITINTSHVEYDTPTR | 601.9672 | Unit | 187.1133 | Unit | 20 | 130 | 16.9 | y3 |
| PgpC (P0AD42) | LQTLQADFVR | 595.8300 | Unit | 949.5101 | Unit | 20 | 130 | 10.12 | y8 |
| PgpC (P0AD42) | LQTLQADFVR | 595.8300 | Unit | 735.3784 | Unit | 20 | 130 | 10.12 | y6 |
| PgpC (P0AD42) | LQTLQADFVR | 595.8300 | Unit | 242.1499 | Unit | 20 | 130 | 10.12 | b2 |
| PgpC (P0AD42) | LQTLQADFVR | 595.8300 | Unit | 343.1975 | Unit | 20 | 130 | 10.12 | b3 |
| PgpC (P0AD42) | DNVTAFPLVQER | 694.8620 | Unit | 959.5308 | Unit | 20 | 130 | 12.04 | y8 |
| PgpC (P0AD42) | DNVTAFPLVQER | 694.8620 | Unit | 888.4937 | Unit | 20 | 130 | 12.04 | y7 |
| PgpC (P0AD42) | DNVTAFPLVQER | 694.8620 | Unit | 741.4253 | Unit | 20 | 130 | 12.04 | y6 |
| PgpC (P0AD42) | DNVTAFPLVQER | 694.8620 | Unit | 230.0771 | Unit | 20 | 130 | 12.04 | b2 |
| PgpC (P0AD42) | VNLIASQIQR | 571.3380 | Unit | 631.3521 | Unit | 20 | 130 | 8.29 | y5 |
| PgpC (P0AD42) | VNLIASQIQR | 571.3380 | Unit | 303.1775 | Unit | 20 | 130 | 8.29 | y2 |
| PgpC (P0AD42) | VNLIASQIQR | 571.3380 | Unit | 214.1186 | Unit | 20 | 130 | 8.29 | b2 |
| PgpC (P0AD42) | VNLIASQIQR | 571.3380 | Unit | 440.2867 | Unit | 20 | 130 | 8.29 | b4 |
| PgpC (P0AD42) | GYGGWVLTMR | 570.2869 | Unit | 919.4818 | Unit | 20 | 130 | 13 | y8 |
| PgpC (P0AD42) | GYGGWVLTMR | 570.2869 | Unit | 520.2911 | Unit | 20 | 130 | 13 | y4 |
| PgpC (P0AD42) | GYGGWVLTMR | 570.2869 | Unit | 407.2071 | Unit | 20 | 130 | 13 | y3 |
| PgpC (P0AD42) | GYGGWVLTMR | 570.2869 | Unit | 221.0920 | Unit | 20 | 130 | 13 | b2 |

**Supplementary Table 6: Mass spectrometry detection settings for phospholipid detection.** Measurements in positive and negative modes were performed separately to circumvent polarity switching. Asterisk (*) indicates species incorporating $^{13}$C-G3P, resulting in a 3 Da mass shift with respect to the regular species (6 Da for PG). Fragmentor voltage and collision energy were adapted from [6]. Novel species were assigned values of similar species. PGP products were not successfully detected. However, the transitions were scanned in all measurements and are therefore included here for completeness. Cell accelerator voltage was 4 V in all conditions.

| Compound class | Compound name | Precursor ion *m/z* | Product ion *m/z* | Fragmentor (V) | Collision energy (eV) | Polarity |
|---|---|---|---|---|---|---|
| Matrix | DOPE | 742.5 | 281.2 | 140 | 25 | Negative |
| Matrix | DOPG | 773.5 | 281.2 | 190 | 37 | Negative |
| Matrix | Cardiolipin | 727.5 | 281.2 | 140 | 29 | Negative |
| Standards | DPPE | 690.5 | 255.1 | 230 | 37 | Negative |
| Standards | DPPG | 721.5 | 255.1 | 220 | 45 | Negative |
| Standards | POPE | 716.5 | 281.2<br>255.1 | 140<br>230 | 25<br>37 | Negative |
| Standards | POPG | 750.5 | 281.2<br>255.1 | 190<br>220 | 37<br>45 | Negative<br>Negative |
| Standards | DOPS | 786.5 | 281.2 | 130 | 40 | Negative |
| 16:0-product | 16:0-LPA* | 412.2 | 155.9 | 150 | 13 | Negative |
| 16:0-product | DPPA* | 650.5 | 255.1 | 210 | 33 | Negative |
| 16:0-product | CDP-DPG* | 955.5 | 712.4 | 135 | 55 | Negative |
| 16:0-product | DPPS* | 737.5 | 255.1 | 180 | 41 | Negative |
| 16:0-product | DPPE* | 693.5 | 255.1 | 230 | 37 | Negative |
| 16:0-product | DPPGP* | 808 | 255.1 | 220 | 45 | Negative |
| 16:0-product | DPPG* | 727.5 | 255.1 | 220 | 45 | Negative |
| 18:1-product | 18:1-LPA* | 438.3 | 155.9 | 160 | 17 | Negative |
| 18:1-product | DOPA* | 702.5 | 281.2 | 190 | 37 | Negative |
| 18:1-product | CDP-DOG* | 1007.5 | 764.5 | 135 | 55 | Negative |
| 18:1-product | DOPS* | 789.5 | 281.2 | 130 | 40 | Negative |
| 18:1-product | DOPE* | 745.5 | 281.2 | 140 | 25 | Negative |
| 18:1-product | DOPGP* | 860 | 281.2 | 190 | 50 | Negative |
| 18:1-product | DOPG* | 779.5 | 281.2 | 190 | 37 | Negative |
| Mixed product | POPA* | 676.5 | 281.2<br>255.1 | 190<br>210 | 37<br>33 | Negative<br>Negative |
| Mixed product | CDP-POG* | 981.5 | 738.5 | 135 | 55 | Negative |

| | | | | | | |
|---|---|---|---|---|---|---|
| Mixed product | POPS* | 763.5 | 281.2<br>255.1 | 130<br>180 | 40<br>41 | Negative<br>Negative |
| Mixed product | POPE* | 719.5 | 281.2<br>255.1 | 140<br>230 | 25<br>37 | Negative<br>Negative |
| Mixed product | POPGP* | 834 | 281.2<br>255.1 | 190<br>220 | 50<br>45 | Negative<br>Negative |
| Mixed product | POPG* | 753.5 | 281.2<br>255.1 | 190<br>220 | 37<br>45 | Negative<br>Negative |

**Supplementary Table 7: Overview of synthesized lipid species.** Symbols between square brackets correspond to residues as defined in the scheme below. Each species is a combination of two acyl chain residues (in numbers) and one head group residue (letter). Lipid species in bold have been quantitatively measured. Species in italic have not been unambiguously detected; however, their presence can be deduced from the successful measurements of subsequent reaction products.

| Enzymes / Precursors | PlsB | PlsC | CdsA | PssA | Psd | PgsA | PgpA |
|---|---|---|---|---|---|---|---|
| Palmitoyl-CoA | 16:0-LPA<br><2,1,a> | DPPA<br><2,2,a> | *CDP-DPG*<br><2,2,b> | DPPS<br><2,2,c> | **DPPE**<br><2,2,d> | *DPPGP*<br><2,2,e> | **DPPG**<br><2,2,f> |
| Oleoyl-CoA | 18:1-LPA<br><3,1,a> | DOPA<br><3,3,a> | *CDP-DOG*<br><3,3,b> | DOPS<br><3,3,c> | **DOPE**<br><3,3,d> | *DOPGP*<br><3,3,e> | **DOPG**<br><3,3,f> |
| Palmitoyl-CoA<br>+<br>Oleoyl-CoA | 16:0-LPA<br><2,1,a><br><br>18:1-LPA<br><3,1,a> | DPPA<br><2,2,a><br><br>DOPA<br><3,3,a><br><br>POPA<br><2,3,a>,<br><3,2,a> | *CDP-DPG*<br><2,2,b><br><br>*CDP-DOG*<br><3,3,b><br><br>*CDP-POG*<br><2,3,b>,<br><3,2,b> | DPPS<br><2,2,c><br><br>DOPS<br><3,3,c><br><br>POPS<br><2,3,c>,<br><3,2,c> | **DPPE**<br><2,2,d><br><br>**DOPE**<br><3,3,d><br><br>**POPE**<br><2,3,d>,<br><3,2,d> | *DPPGP*<br><2,2,e><br><br>*DOPGP*<br><3,3,e><br><br>*POPGP*<br><2,3,e>,<br><3,2,e> | **DPPG**<br><2,2,f><br><br>**DOPG**<br><3,3,f><br><br>**POPG**<br><2,3,f>,<br><3,2,f> |
| Palmitoyl-CoA<br>+<br>NBD-palmitoyl-CoA | 16:0-LPA<br><2,1,a><br><br>*NBD-LPA*<br><4,1,a> | DPPA<br><2,2,a><br><br>NBD-PPA<br><2,4,a>,<br><4,2,a><br><br>*NBD-NDB-PA*<br><4,4,a> | *CDP-DPG*<br><2,2,b><br><br>*CDP-NBD-PG*<br><2,4,b>,<br><4,2,b><br><br>*CDP-NBD-NBD-G*<br><4,4,b> | DPPS<br><2,2,c><br><br>NBD-PPS<br><2,4,c>,<br><4,2,c><br><br>*NBD-NDB-PS*<br><4,4,c> | **DPPE**<br><2,2,d><br><br>NBD-PPE<br><2,4,d>,<br><4,2,d><br><br>*NBD-NDB-PA*<br><4,4,d> | - | - |

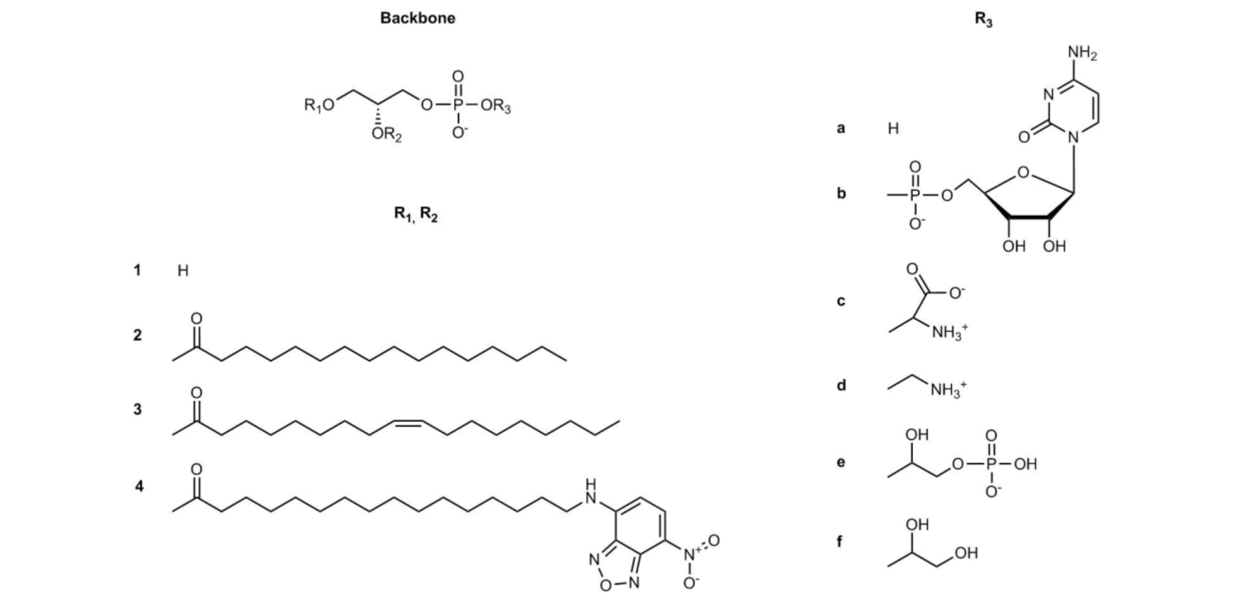

### SUPPLEMENTARY FIGURES

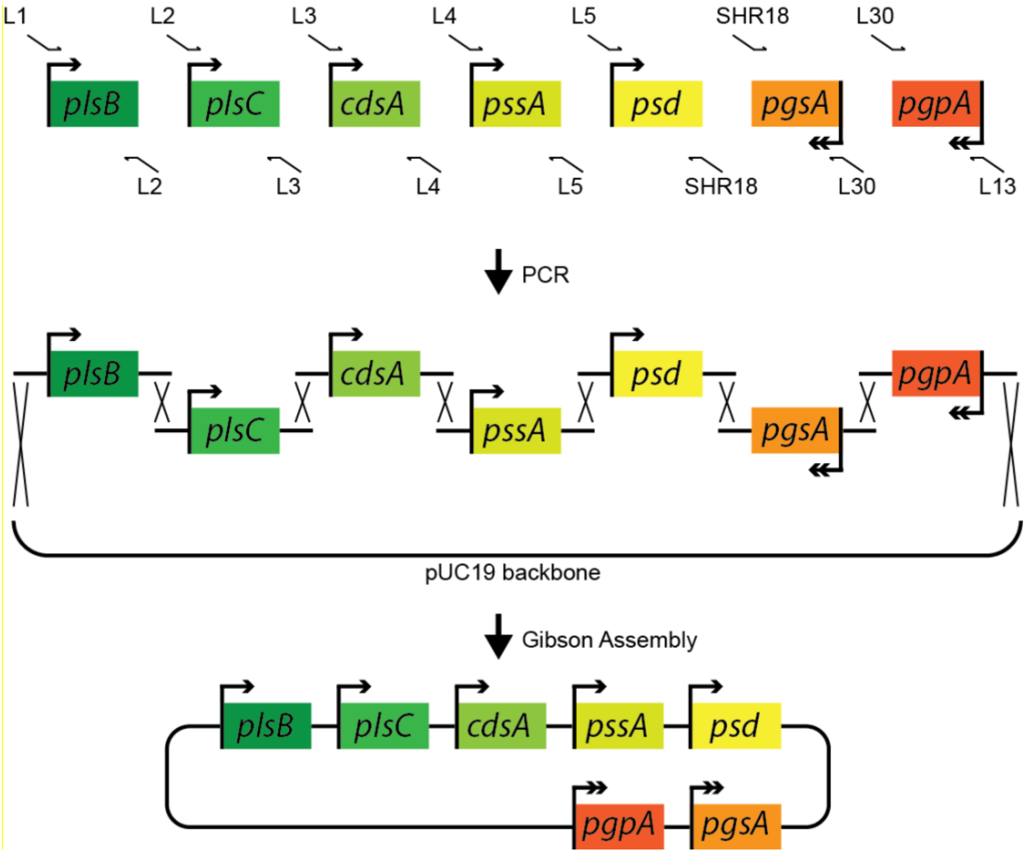


**Supplementary Figure 1: Scheme for the construction of the pGEMM7 plasmid.** Homologous linker sites are added to the transcriptional cassettes via PCR in the first step. After purification of the PCR fragments the entire plasmid is assembled by Gibson assembly in a single reaction.

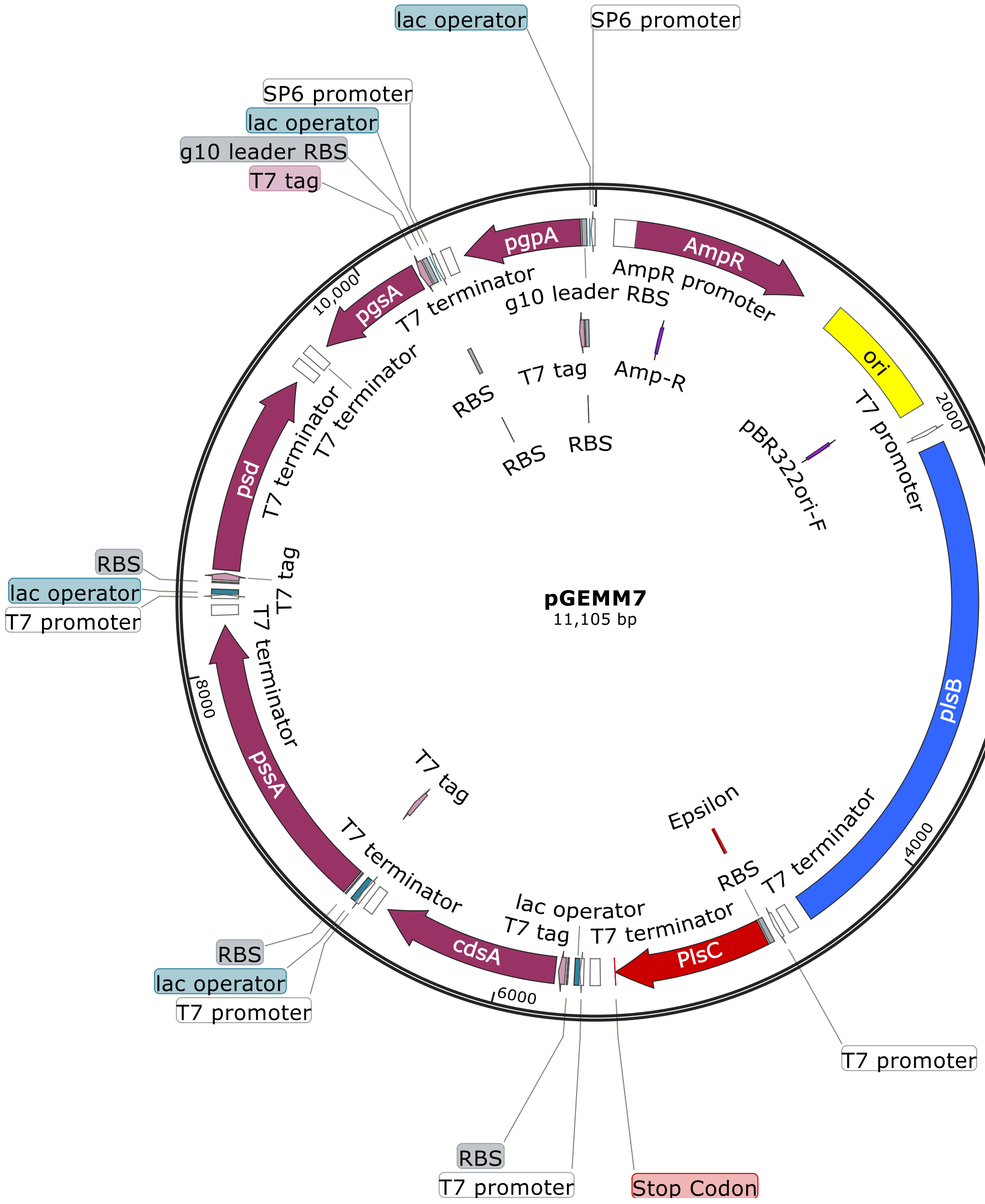


**Supplementary Figure 2: Sequence map of the pGEMM7 construct.** The plasmid map was generated with Snapgene.

**a**

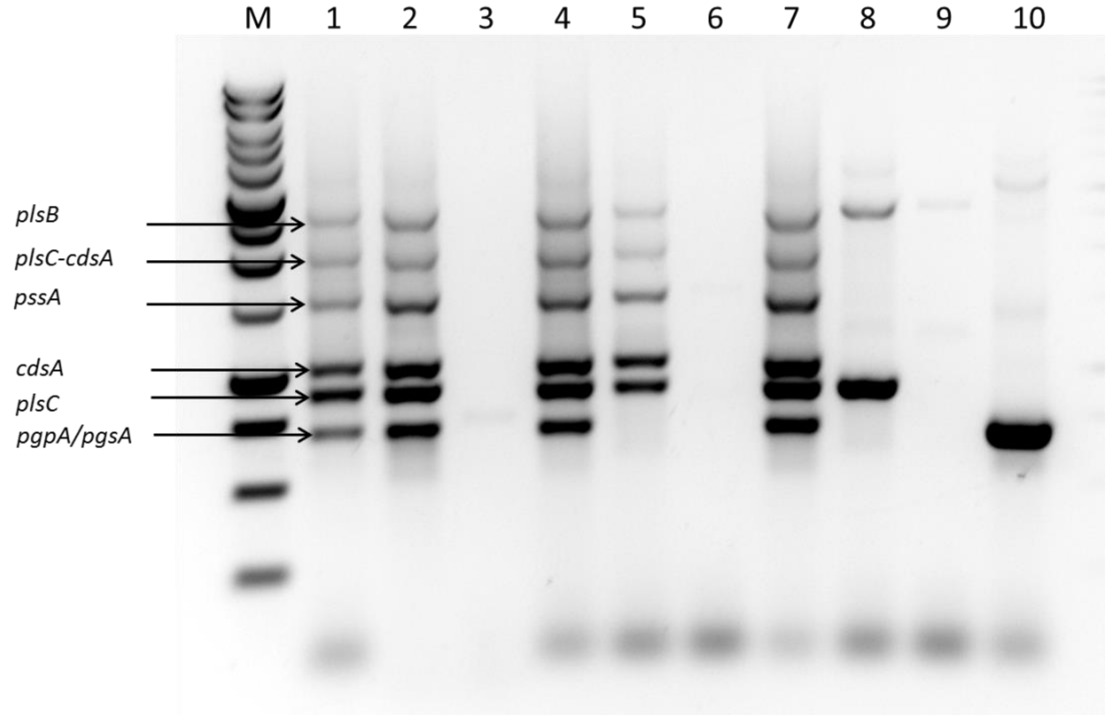


**b**

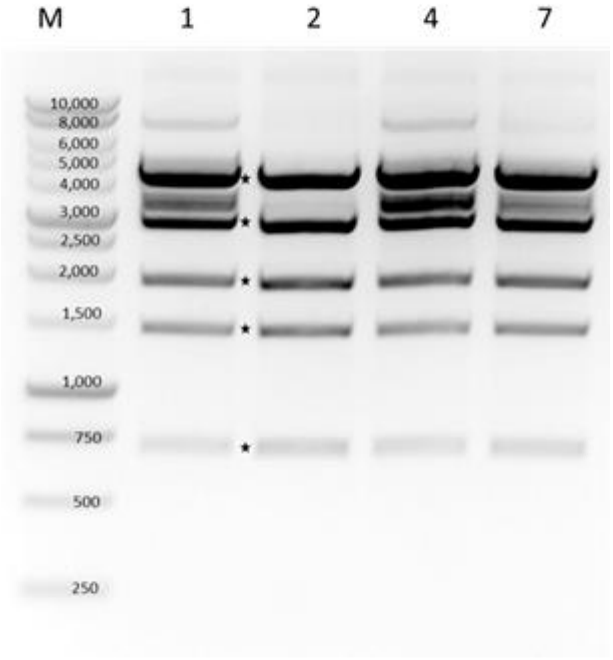


**Supplementary Figure 3: Verification of pGEMM7 construct. a**, Agarose gel of a colony PCR using primers 91 and 397. Colonies 1, 2, 4 and 7 were chosen to be further analysed due to the observed band pattern. **b**, Agarose gel stained with SYBR Safe DNA dye of a restriction digestion of pGEMM7. 1 µg of total DNA was loaded. 5 µL of Benchtop 1kb ladder (Promega) were loaded in the first lane. The stars indicate the location of the expected bands. The numbers above the lanes indicate the colony number that was tested.

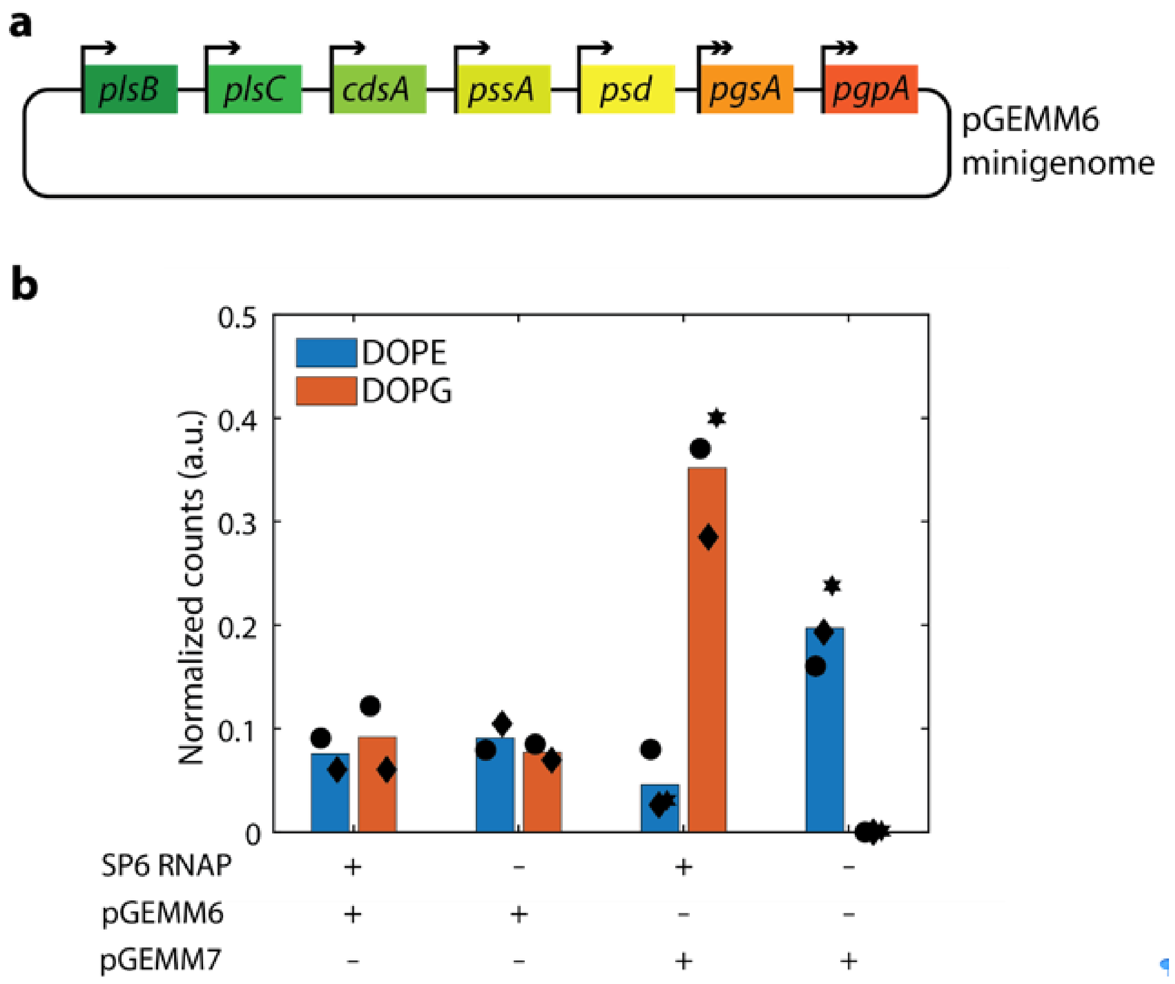


**Supplementary Figure 4: Read-through causes non-specific PG synthesis with pGEMM6. a**, Schematic representation of the pGEMM6 plasmid where all five genes under T7 promoter control (singel arrow) were assembled in the same orientation as the two genes *pgsA* and *pgpA* under SP6 promoter control (depicted by the double arrow). **b**, Synthesis of phospholipids from pGEMM6 (data with pGEMM7 are appended for comparison), in the presence of LUVs and all necessary precursors, with and without SP6 RNAP. In the absence of SP6 RNAP, *pgsA* and *pgpA* should not be expressed, and therefore, no DOPG should be synthesized. However, an appreciable yield of DOPG was observed without SP6 RNAP in the case of pGEMM6. We hypothesized that this was caused by transcription termination read-through by the T7 RNAP, transcribing downstream PG synthesis genes even if those are under control of an orthogonal promoter. Bars are average values from two independent repeats (three with pGEMM7), each represented by a different symbol.

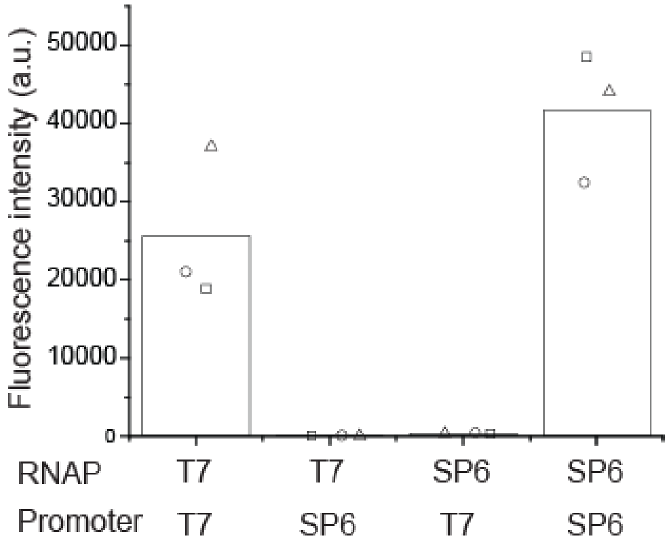


**Supplementary Figure 5: Orthogonality diagram of T7 vs Sp6 promoters.** The pUC57-T7p-LacO-meYFP-LL-spinach-T7t and pUC57-SP6p-LacO-meYFP-LL-spinach-T7t were expressed in an *E. coli* cell lysate in the presence of either the T7 or SP6 RNAP. End-point fluorescence signal from YFP was only observed when the RNAP matched with its corresponding promoter, indicating strong orthogonality of both promoter/RNAP pairs. Bars represent mean values from three repeats and the different symbols are individual data points.

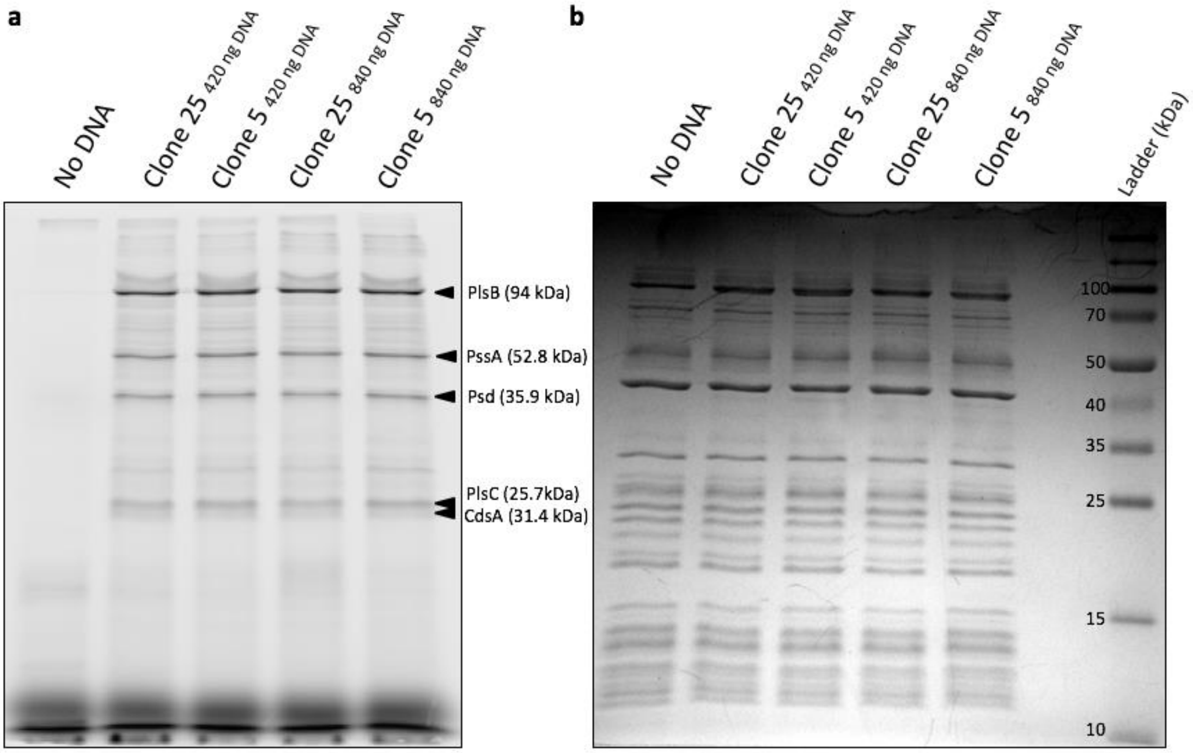


**Supplementary Figure 6: Co-translational labelling of proteins with GreenLys.** Early versions of pGEMM6 and pGEMM7 were screened for their ability to produce lipid-synthesizing enzymes. Clones 5 and 25 encode five genes that were successfully expressed in PURE*frex*2.0 supplemented with 1 µL of GreenLys reagent (Promega). The translation products were analysed by SDS-PAGE and Typhoon imaging. **a**, Fluorescence image with appended arrows pointing to the relevant protein bands. The protein names and theoretical molecular masses are indicated. Note that CdsA migrates significantly lower than predicted in the gel, as previously reported [6]. **b**, CBB stain image of the gel shown in a. Background proteins from the PURE system are visible, whereas the yield of synthesized proteins is too low to be directly observed.

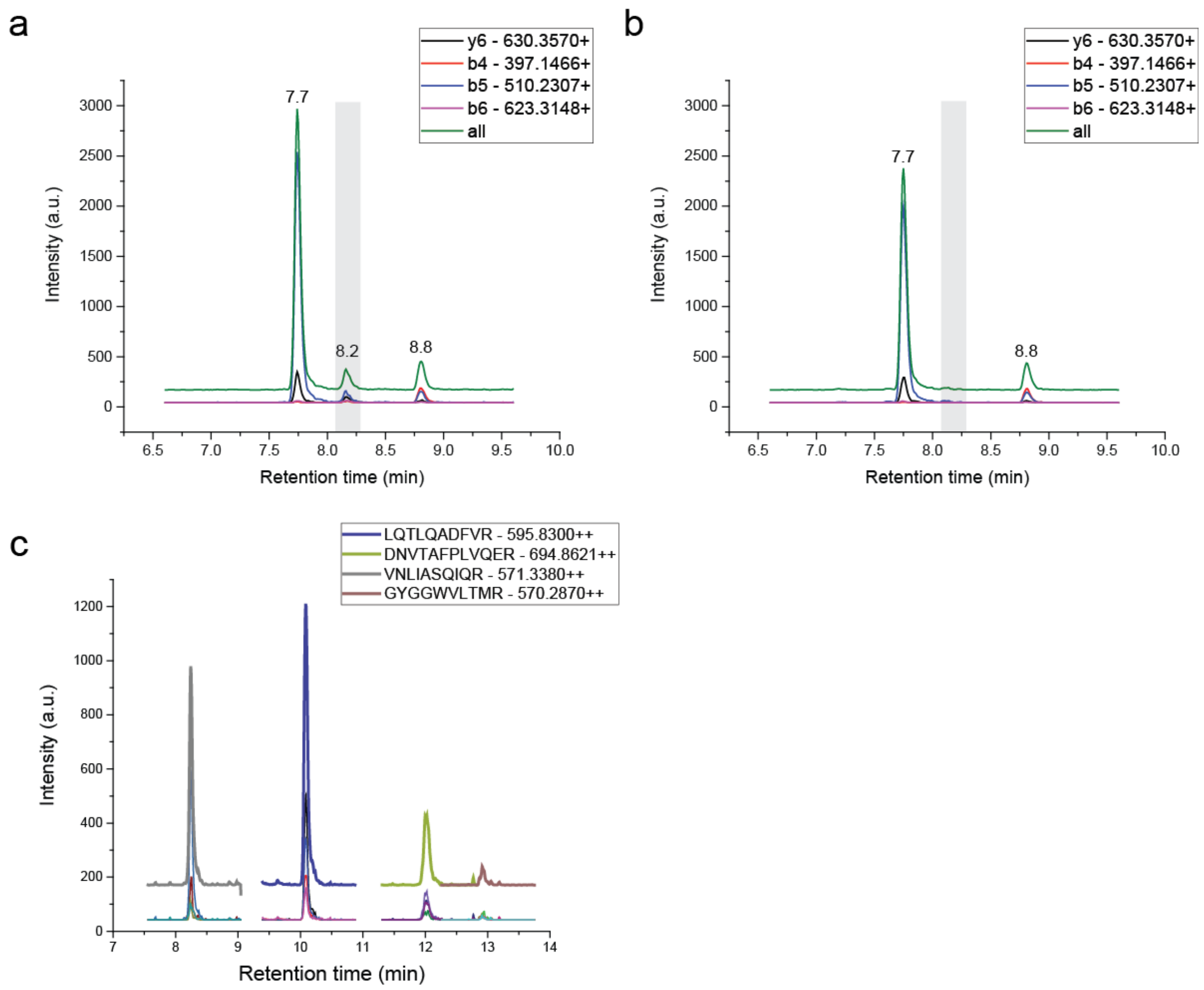


**Supplementary Figure 7: LC-MS detection of CdsA by a single specific peptide DSGHLIPGHGGILDR and PgpC by four peptides.** Peaks corresponding to tryptic peptides of the other enzymes were clearly identified and the chromatograms are not shown here. Their specifications can be found in **Supplementary Table 4**. **a**, Plotted intensities of four fragments of the CdsA peptide together with the combined intensities of all fragments. The overlayed gray area indicates the expected position at 8.2 min of the specific peak in the chromatogram. This specific peak is absent in a control cell-free protein expression reaction incubated without DNA (**b**). The visible peaks at 7.7 minutes and 8.8 min retention time can be attributed as background signals from the PURE*frex*2.0 proteins themselves or other contaminants. **c**, Chromatograms of all transitions and the combined measured ion current of each peptide of PgpC that were detectable after trypsin digestion. Source Data are available for panels a and b.

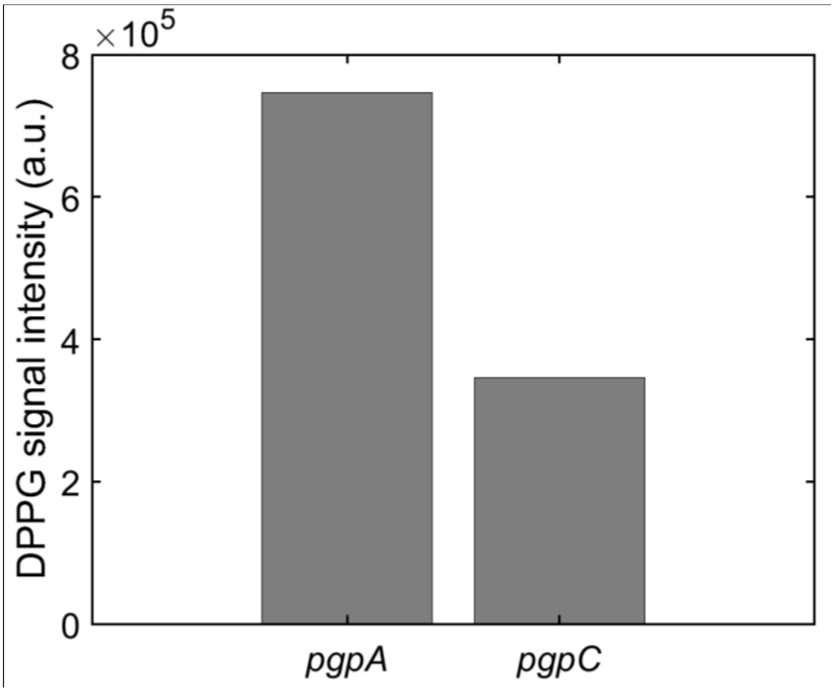


**Supplementary Figure 8: PgpC can replace PgpA to synthesize PG.** PURE*frex*2.0 expression of *plsB*, *plsC*, *cdsA*, *pgsA*, and *pgpA* or *pgpC* (10 nM each), in the presence of LUVs, palmitoyl-CoA, and all necessary lipid synthesis precursors. Genes were added as separate DNA molecules. Replacing PgpA by PgpC results in lower but appreciable yield of PG. Experiment was performed once.

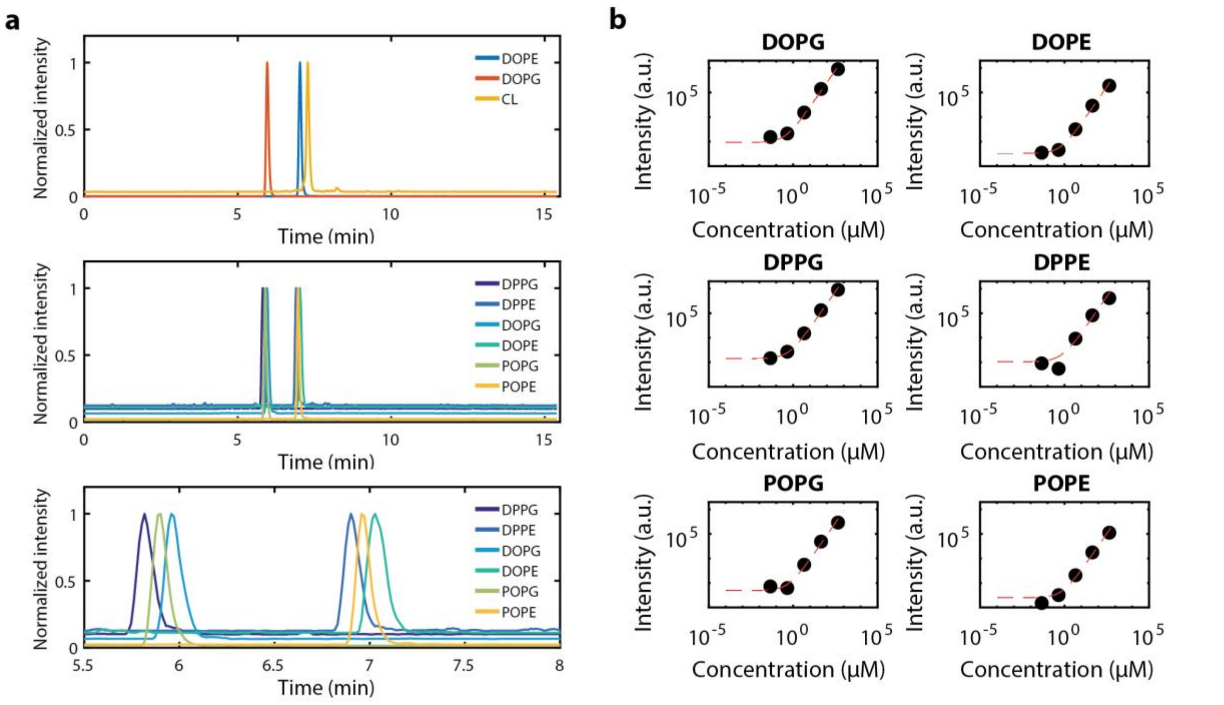


**Supplementary Figure 9: LC-MS detection of phospholipids. a**, Representative LC chromatograms of the studied phospholipids. Experimental conditions are as described in **Fig. 1c**. Two AU of SP6 RNAP, along with a 1:1 mix of oleoyl- and palmitoyl-CoA were supplemented to LUVs. Peak intensity was normalized to aid visualization. (top) Chromatograms of the lipids present in the liposome matrix. (middle, bottom) Chromatograms of synthesized phospholipids. The lower panel is a zoom-in graph, showing the typical elution pattern. Species with longer (oleoyl) chains elute later than species with shorter (palmitoyl) chains. **b**, Representative calibration curves for quantification of phospholipid concentrations. Five different known concentrations of the indicated phospholipids were injected, from low to high concentration, and plotted versus the resulting total integrated counts (TIC) of the corresponding peak labelled as 'Intensity' on the graphs. A linear fit was computed to obtain a calibration curve for each lipid (overlaid red dashed line). TIC values from cell-free synthesized lipids were then converted into absolute concentrations. Source Data are available for panel b.

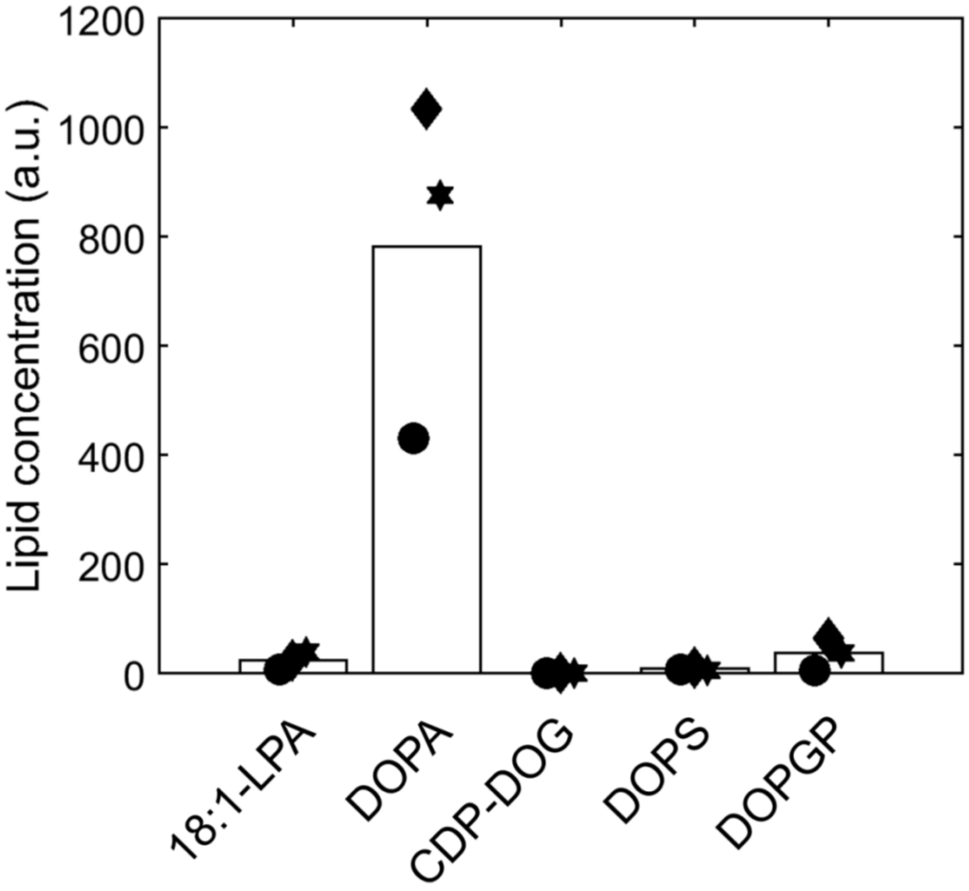


**Supplementary Figure 10: Accumulation of lipid synthesis intermediates.** Signal corresponding to all five lipid synthesis intermediate species. The measurements correspond to the rightmost data points shown in **Fig. 1c**, that is expression of pGEMM7 in the presence of LUVs and 4 AU of SP6 RNAP. Only DOPA accumulates in significant amounts. Other species have a total integrated peak intensity below 100 a.u.; therefore, the signals cannot be distinguished from noise. Source Data are available.

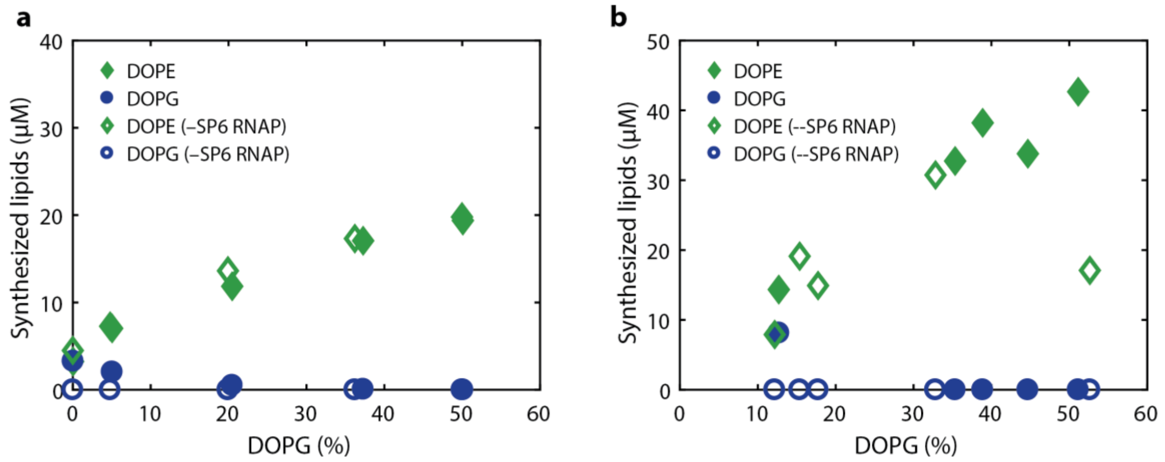


**Supplementary Figure 11: Two repeats of the experiments shown in main text Fig. 2b. a** is an identical repeat. In **b** other DOPG membrane concentrations where chosen. Source Data are available.

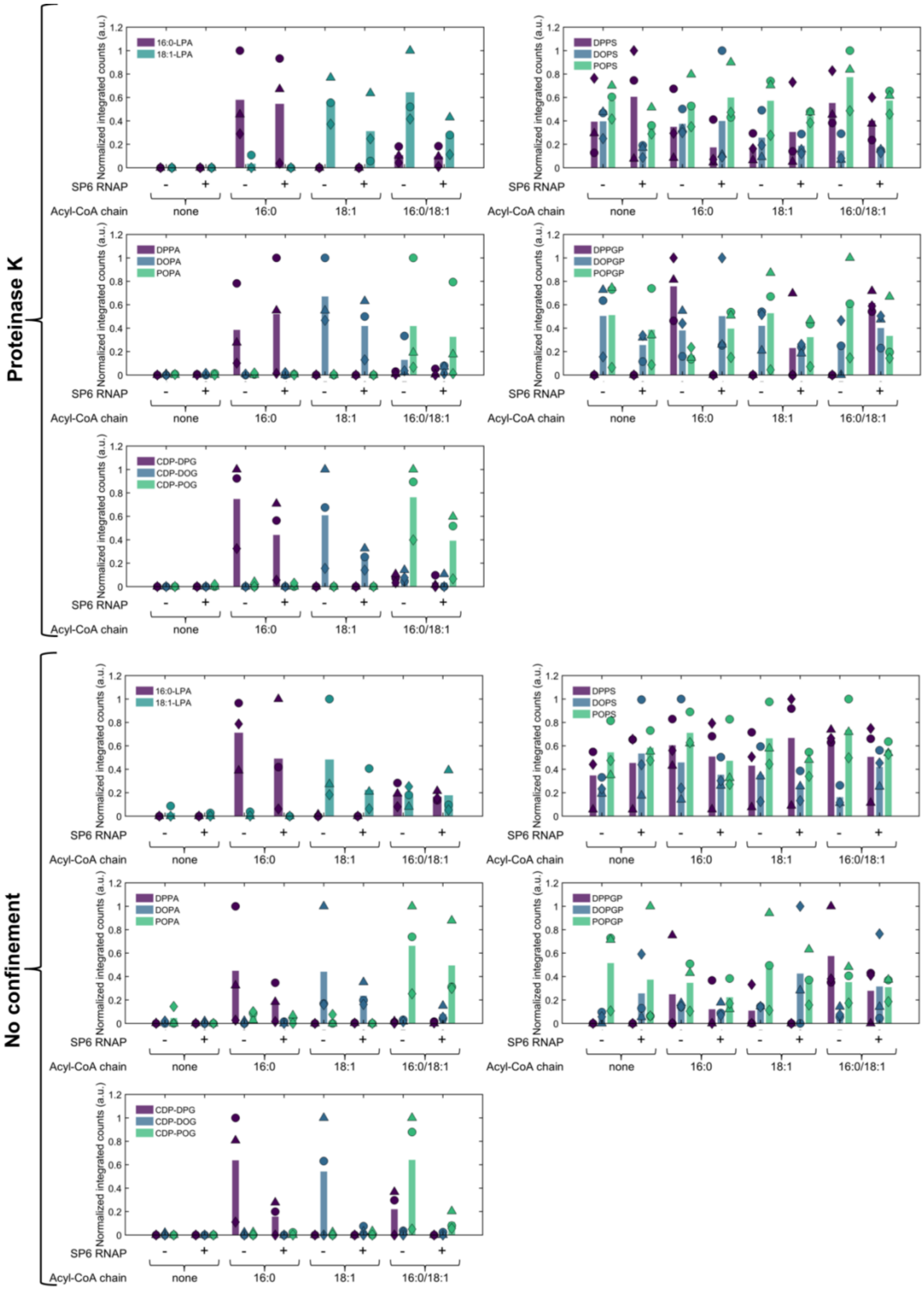


**Supplementary Figure 12: LC-MS detection of synthesized phospholipids and accumulated intermediate products in giant liposome experiments.** Phospholipid intermediates LPA, PA, and CDP-DAG, but not PGP and PS, were detected. Data are plotted as total integrated counts normalized to the highest value per species, for all intermediate lipid species. Dataset was obtained from the same samples as shown in main text **Fig. 3d,e**. Bars indicate mean values. Symbols indicate values from individual experiments and correspond to data shown main text **Fig. 3d,e**. Precursors palmitoyl-CoA (16:0 acyl-CoA) and oleoyl-CoA (18:1 acyl-CoA) were used as indicated. Addition of proteinase K in the external medium confines gene expression to the interior of liposomes (upper panels). For LPAs, PAs and CDP-DAGs, signals were observed as expected, i.e. di-palmitoyl (DP) or di-oleoyl (DO) products when starting with 16:0 or 18:1 acyl-CoA, respectively, and both DP, DO and mixed chain products (PO) when starting with a mixture of the two precurscors. For PS and PGP species, no clear pattern was detected and only very low peak intensities (< 100 integrated counts) were measured. Because successful detection of PS was confirmed in other experiments (**Supplementary Fig. 16**), the results indicate that PS species do not accumulate under these conditions. In contrast, PGP signal was never observed, suggesting that its detection requires optimized protocols. Accumulation of LPAs, PAs, and CDP-DAGs was lower in the presence of SP6 RNAP when the PG synthesis pathway is active. Presumably, having two downstream pathways instead of one might result in a higher flux through the upstream (pre-branchpoint) part of the pathway. Source Data are available.

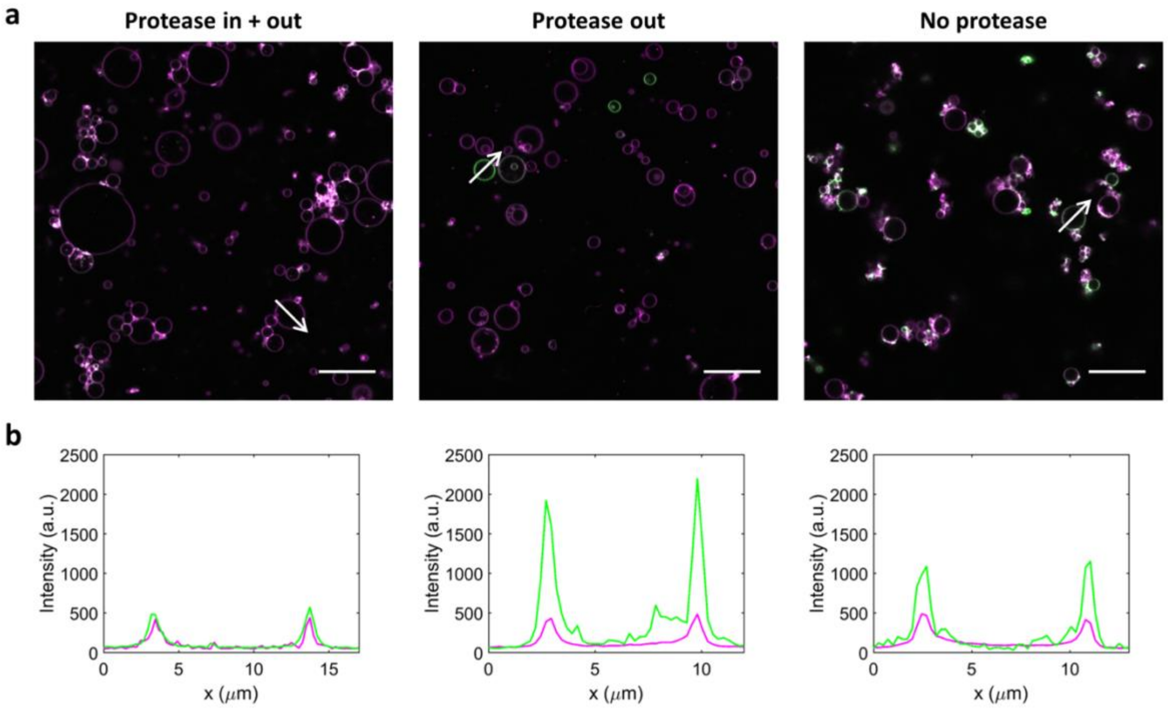


**Supplementary Figure 13: Confocal fluorescence images of giant liposomes with synthesized NBD-labelled phospholipids.** Experimental conditions are as indicated in main text **Fig. 4d**. Briefly, liposomes encapsulating pGEMM7 and PURE*frex*2.0 were prepared and incubated in the presence of NBD-palmitoyl-CoA and palmitoyl-CoA (1:9 mol. fraction). **a**, Representative images of different liposome samples are shown. Proteinase K was co-encapsulated inside liposomes, preventing gene expression and thus lipid synthesis (left). Proteinase K was added to preformed liposomes, preventing lipid synthesis from occurring outside the liposome lumina (middle). When no proteinase K was supplemented, gene expression and lipid synthesis can take place both inside and outside liposomes (right). NBD fluorescence is shown in green and the Texas Red-dyed liposome membrane in magenta. Scale bars represent 20 µm. **b**, Line profiles of Texas Red (red) and NBD (green) signals along the arrows appended in **a**. a.u., arbitrary unit.

a

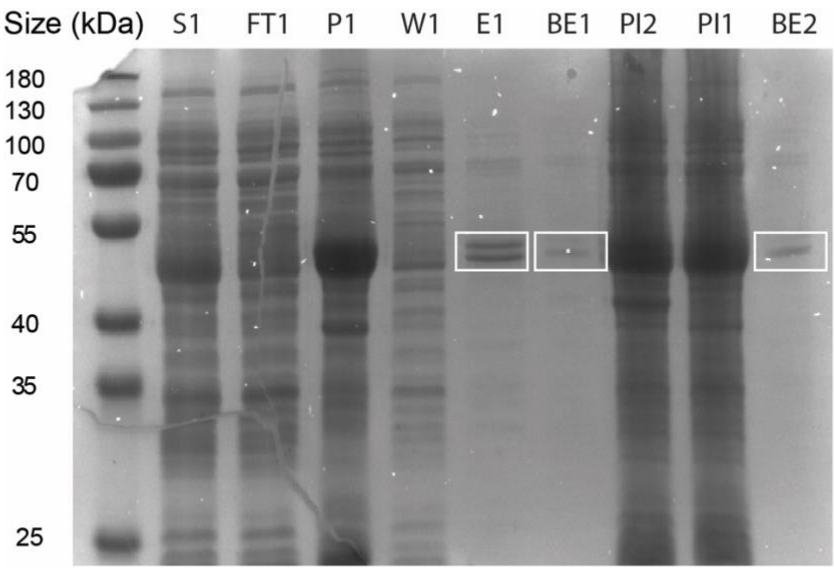


b

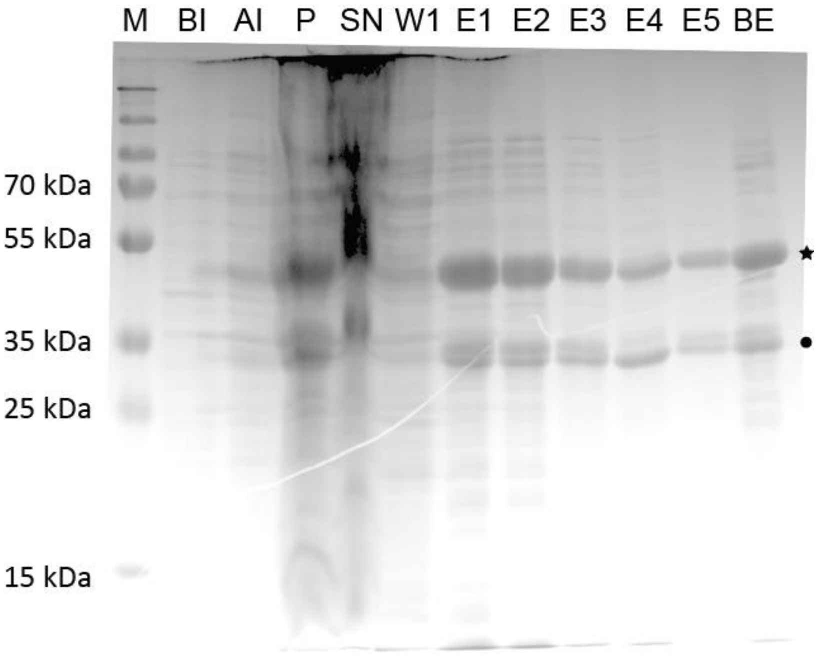


**Supplementary Figure 14: Purification of LactC2-eGFP and LactC2-mCherry.** Images of SDS-PAGE gels to visualize the purification steps of LactC2-eGFP (**a**) and LactC2-mCherry (**b**). **a**, Recombinant LactC2-eGFP was expressed in *E. coli* and purified using Ni-NTA chromatography column. "1" stands for the protein produced by strain Rosetta ER2566, "2" by strain Rosetta 2; "PI" stands for post-induction with IPTG, "S" for the supernatant and "P" for the pellet, both after cell lysis by sonication, "FT" for the flow-through of the membrane binding step, "W" for the flow-through of the column washing step, "E" for eluate and "BE" for the final purified protein after buffer exchange. The white boxes indicate the expected region for the protein band. Double bands occurring in lane E1 can hint at incomplete reduction of the protein. This is supported by the fact that after buffer exchange there is only a single band visible which could indicate fresh and sufficient amounts of DTT fully reduced all disulfide bonds. **b**, In lane “M” 5 µL of Page ruler plus ladder (Promega) was loaded. In the following lanes a sample of the expression strain before induction (BI), after overnight induction at RT (AI), the pellet after lysis and centrifugation (P), the supernatant after lysis (SN), the first washing step (W1), elutions 1 to 5 (E1-E5) and a sample after buffer exchange (BE). The expected product size of the fusion protein is 47 kDa corresponding to the bands marked by a star. The dot marker indicates a set of two bands roughly corresponding to a monomeric mCherry protein including the T7 tag (30 kDa).

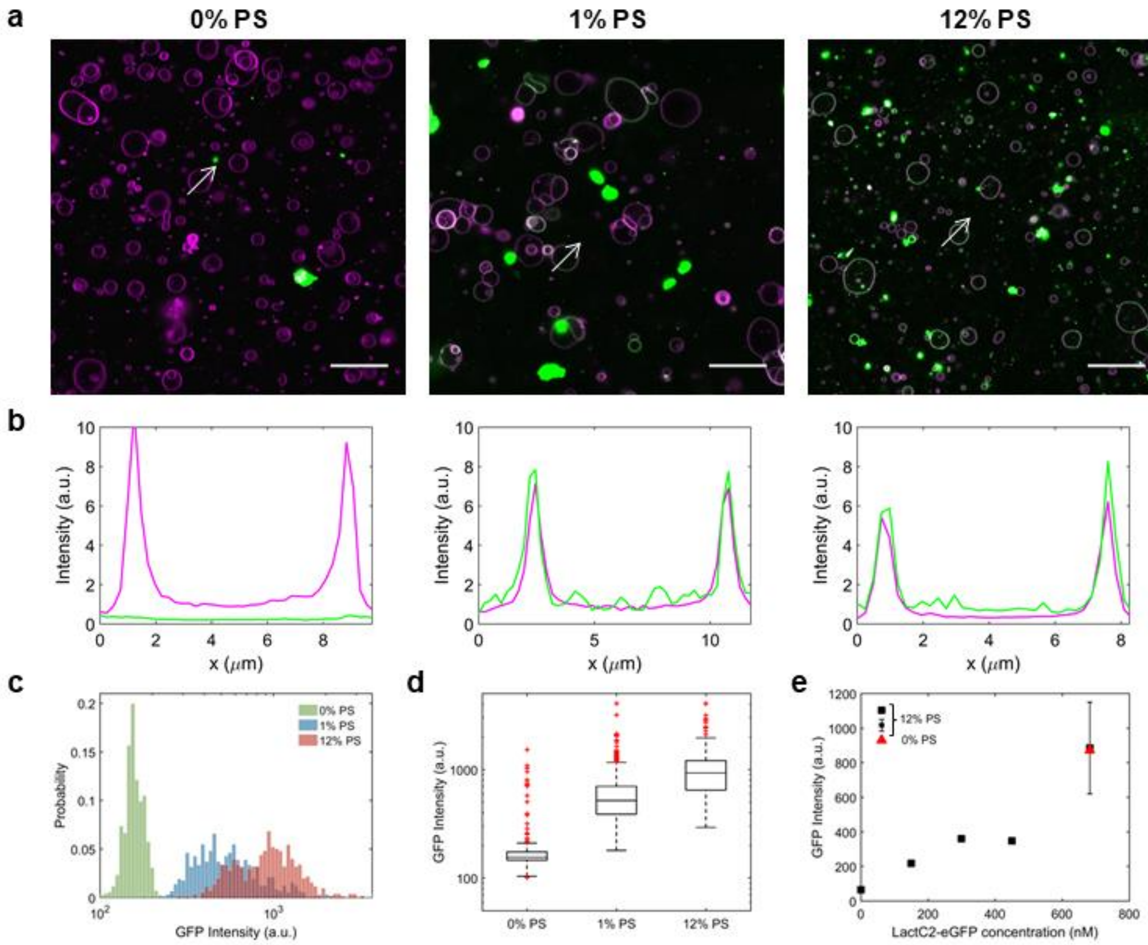


**Supplementary Figure 15: LactC2-eGFP specifically binds to PS-containing liposomes. a** Confocal fluorescence images of liposomes (membrane in magenta) containing 0%, 1%, or 12% DOPS in the membrane. The liposomes were incubated with 150 nM of LactC2-eGFP (green signal) before imaging. Green spots are LactC2-eGFP aggregates, which were typically observed in all conditions and do not colocalize with vesicles. Scale bars represent 20 µm. **b**, Fluorescence intensity line profiles corresponding to the arrows appended in **a**. Texas Red membrane dye signal is in magenta and LactC2-eGFP signal is in green. Clear signal colocalization can be seen in the presence of 1% and 12% PS. **c**, Average LactC2-eGFP rim intensity for 0%, 1% and 12% PS. Number of analyzed liposomes *N* = 675, 560, 496, respectively. **d,** Box plot of the data displayed in **c**. *N* = 675, 560, 496 (from left to right). The box represents all data point between the 25$^{th}$ and 75$^{th}$ percentile, with the line indicating the median. Whiskers extend the most extreme data points not considered outliers. Outliers are plotted individually as red '+' signs. **e**, Average LactC2-eGFP rim intensity for liposomes with 0% or 12% PS and varying concentrations of LactC2-eGFP. For 12% PS, *N* = 503, 273, 93 and 403 liposomes (from left to right), all from one experiment. The error bar corresponds to the mean ± SD of two 12% PS samples, with *N* = 314 and *N* = 71. The red triangle represents the average GFP intensity of 60 liposomes, from one experiment, for liposomes containing 0% PS. Clear non-specific binding of the LactC2-eGFP probe to the membrane can be observed at high concentration. To ensure PS-binding specificity, a working concentration of LactC2-eGFP of 150 nM was employed in all measurements with expressed pGEMM7. a.u., arbitrary unit. Source Data are available for panels c-e.

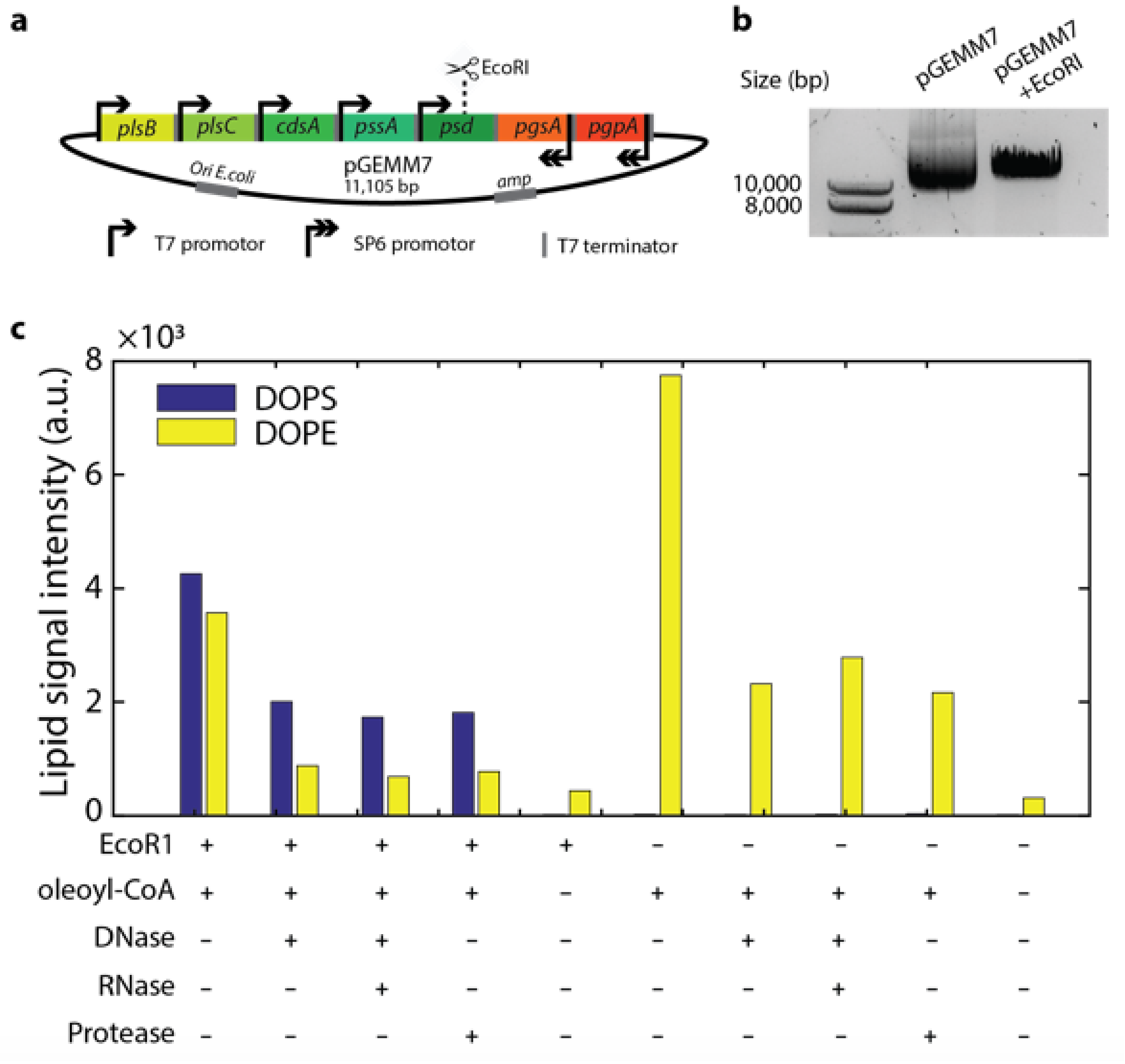


**Supplementary Figure 16: Expression of EcoRI-restricted pGEMM7 results in accumulation of phosphatidylserine. a**, Schematic representation of the pGEMM7 plasmid map. The EcoRI site in the *psd* gene is depicted. **b**, 1% agarose gel showing unrestricted pGEMM7 and pGEMM7 cleaved with EcoRI. **c**, LC-MS analysis of synthesized DOPS and DOPE with circular pGEMM7 (–EcoRI) or linearized pGEMM7 (+EcoRI) expressed inside giant liposomes. No SP6 RNAP was added. The total integrated counts for DOPS and DOPE are shown. Accumulation of PS is only observed with the linear DNA template, i.e. when expression of the Psd enzyme which converts PS into PE is inhibited. Some residual PE synthesis is observed, probably due to incomplete enzymatic digestion of pGEMM7. Various methods to confine gene expression to the inside of vesicles have been tested. Using proteinase K or DNase gave equivalent results, corroborating the findings presented in the main text **Fig. 3a**. Source Data are available for panel c.

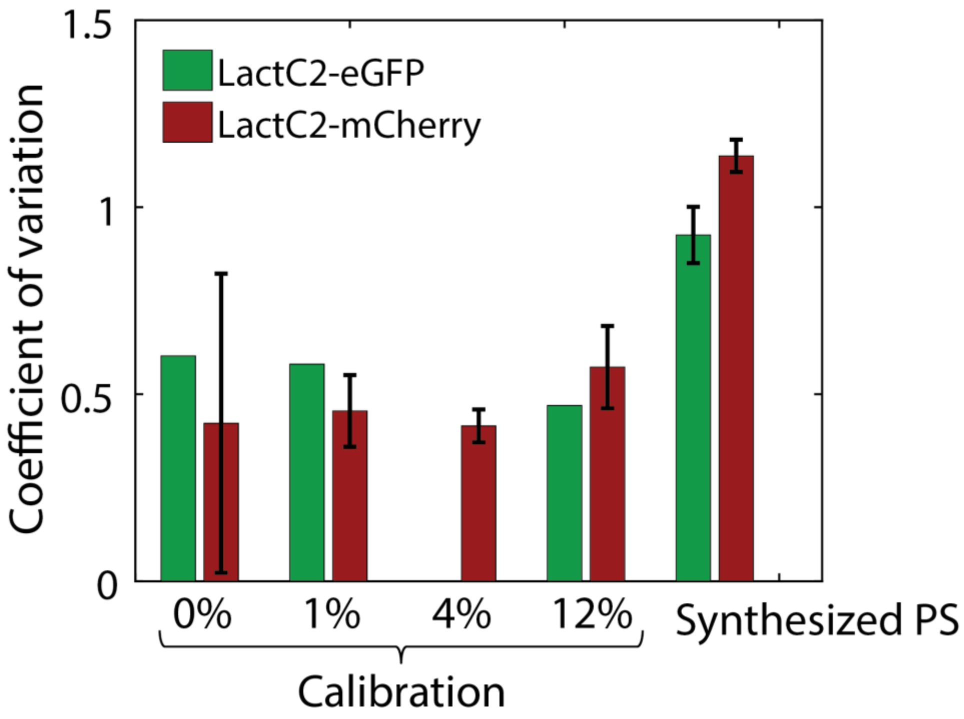


**Supplementary Figure 17: Liposome-compartmentalized PS synthesis results in a higher coefficient of variation in LactC2-eGFP signal than when PS is directly included in the vesicle membrane.** The coefficient of variation, defined as $\sigma/\mu$, where $\sigma$ the standard deviation and $\mu$ the mean, of LactC2-eGFP or -mCherry was calculated for control samples with liposomes containing a fixed amount of PS (0, 1, 4 or 12 mol. %, as indicated) and for in-liposome synthesized PS samples. Liposome-confined synthesis of PS results in higher values of the coefficient of variation with both fluorescent probes, providing a quantitative measure of the liposome-to-liposome heterogeneity in the yield of synthesized PS. When an error bar is displayed, data represents the mean ± SD of two different biological samples, except for synthesized PS with LactC2-eGFP, where the mean ± SD of three biological replicates is shown. No error bar indicates that the experiment was performed once. Source Data are available.

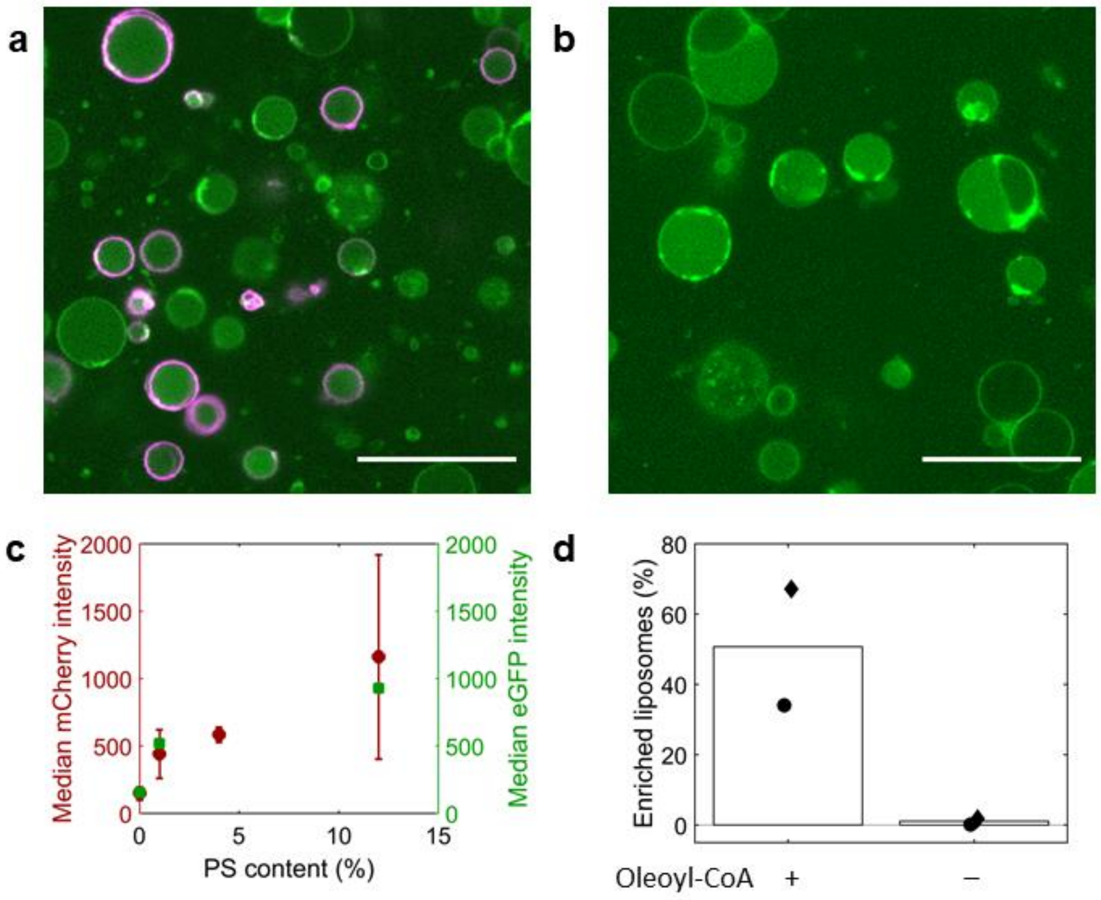


**Supplementary Figure 18: LactC2-mCherry is also a compatible PS-specific binding probe.** Confocal fluorescence images of liposomes encapsulating all necessary components for pGEMM7Δ*psd*-directed lipid synthesis, in the presence (**a**) or absence (**b**) of oleoyl-CoA. LactC2-mCherry signal is displayed in magenta. Since mCherry is not spectrally compatible with the Texas Red membrane dye, liposomes were alternatively stained by addition of acridine orange (AO, green). AO is commonly used as a nucleid acid dye, but we have previously observed that the hydrophobic molecule partitions also into liposome membranes [7]. Compared with the eGFP fusion protein, less prominent clusters of LactC2-mCherry were observed. Scale bar represents 20 µm. **c**, Median LactC2-mCherry intensity values for liposomes containing various amounts of PS (indicated in mol. %). Data are the mean ± SD from three repeats. Data obtained with LactC2-eGFP (**Supplementary Fig. 15**) are superimposed for comparison. **d**, Percentage of PS-enriched liposomes after expression of pGEMM7Δ*psd* in the presence or absence of oleoyl-CoA. Bars represent the mean from two independent repeats. Individual data points are marked with a different symbol. Results are similar as with LactC2-eGFP (main text **Fig. 5e**) demonstrating that the nature of the fluorescent protein does not influence the membrane binding ability, despite the fact that LactC2-eGFP is more prone to aggregation. Source Data are available for panels c and d.

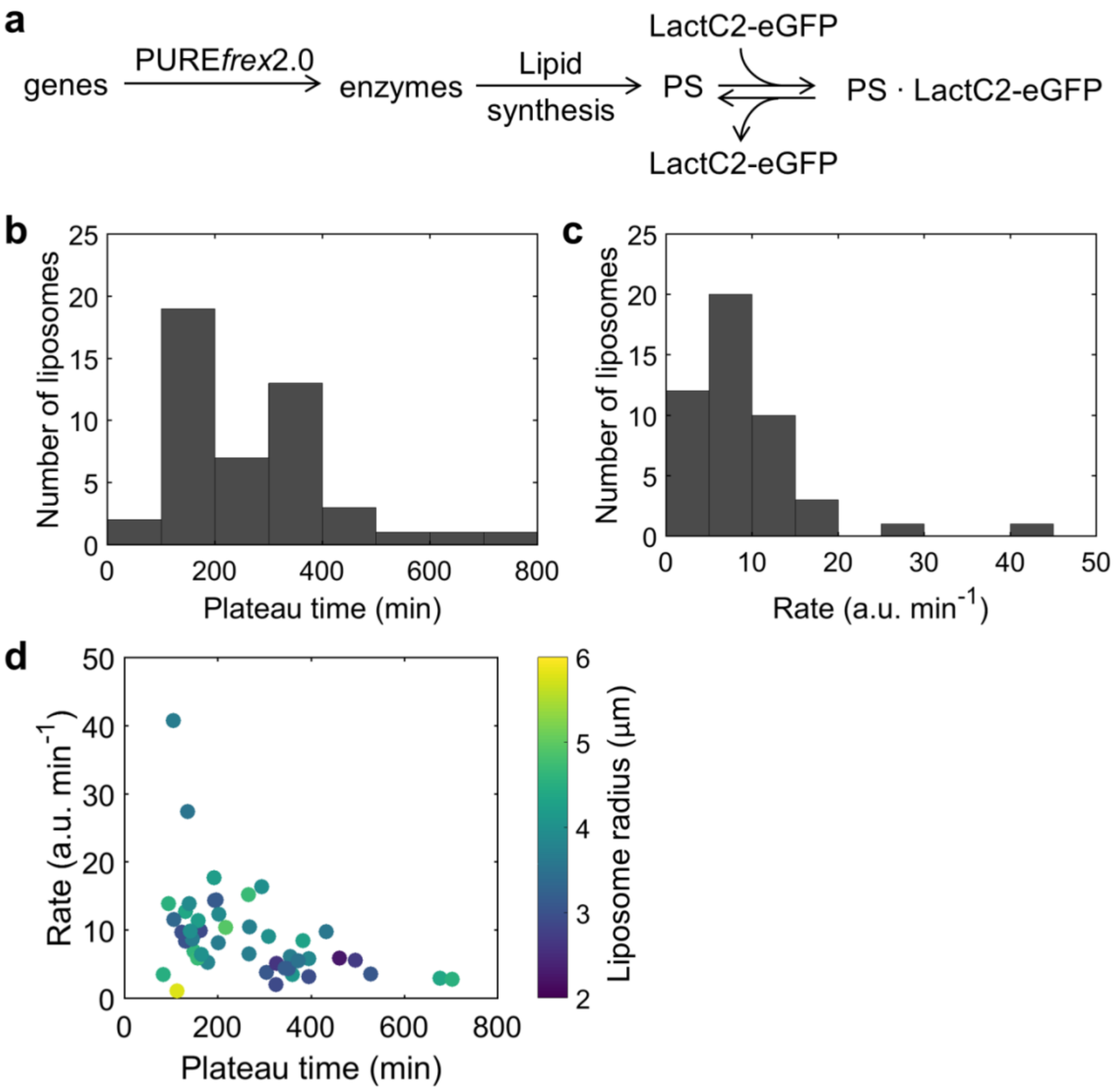


**Supplementary Figure 19: Kinetic analysis of combined enzyme expression, PS synthesis, and LactC2-eGFP binding.** **a**, Overview of the main kinetic steps resulting in binding of LactC2-eGFP to synthesized PS, as observed in main text **Fig. 5**. **b**, **c**, Histograms, for the 47 traces shown in **Fig. 5g**, of plateau time (**b**) and apparent rate (**c**), as defined in Eq. 2 and Eq. 3, respectively. **d**, Scatter plot of the plateau time and apparent rate as shown in **b** and **c**. Color coding corresponds to the (apparent) liposome radius [4]. No clear size dependence can be observed. Plateau time and rate are somewhat negatively correlated (Pearson's correlation coefficient ρ = –0.44). Source Data are available for panels b-d.

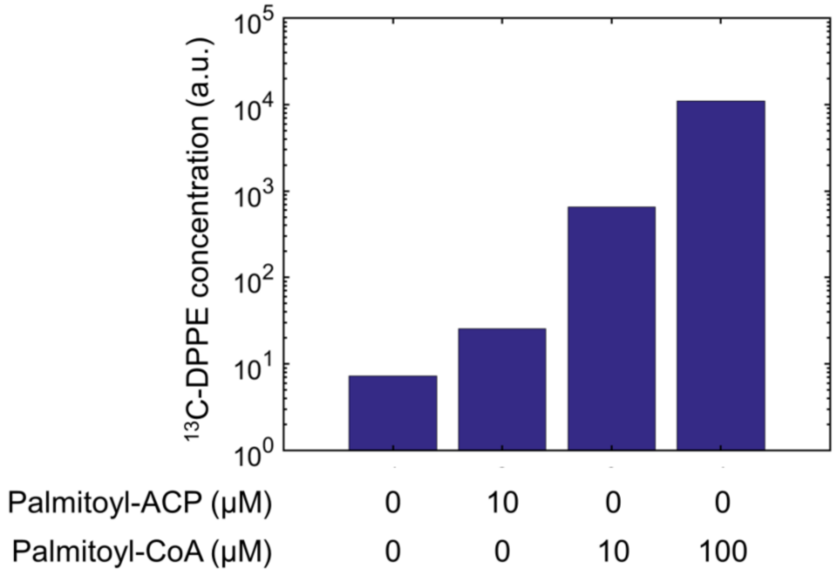


**Supplementary Figure 20: Palmitoyl-ACP is an inefficient acyl chain donor for pGEMM7-directed lipid synthesis.** DPPE was synthesized by expressing pGEMM7 in the presence of LUVs and all necessary precursors. As acyl-chain donor, palmitoyl-acyl carrier protein (palmitoyl-ACP) and palmitoyl-CoA were used. Palmitoyl-ACP was taken from a stock solution of 1.65 mg mL$^{-1}$ in 10 mM $MgSO_4$. Biosynthesis yield of DPPE from palmitoyl-ACP (10 µM) was more than one order of magnitude lower than the yield using an equal concentration of palmitoyl-CoA. Final concentration of input palmitoyl-ACP was limited by the stock concentration. In contrast, palmitoyl-CoA could be used up to 100 µM, leading to a yield of synthesized DPPE that is over two orders of magnitude higher compared to that with 10 µM palmitoyl-ACP. These results, combined with the fact that using acyl-ACP introduces a cumbersome extra protein purification step, made us decide to not further investigate acyl-ACP as a phospholipid precursor. Y-axis is total integrated count. Experiment was performed once.

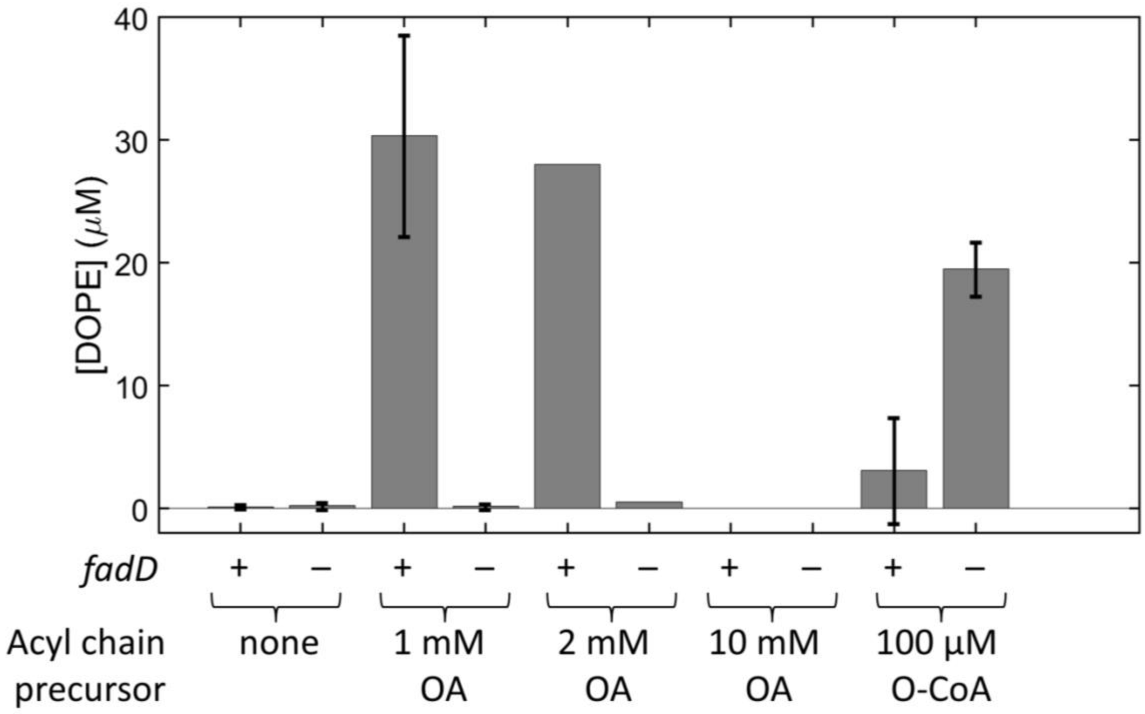


**Supplementary Figure 21: Oleic acid can be utilized as a phospholipid precursor by co-expressing pGEMM7 and the *fadD* gene.** Lipid synthesis enzymes encoded by pGEMM7 were produced in the presence of LUVs and various amounts of oleic acid (OA) or oleoyl-CoA (O-CoA), with or without *fadD* DNA. In all conditions, 50 μM CoA was added. Data are absolute concentrations of synthesized DOPE as measured by LC-MS. Single or two independent experiments were conducted, as indicated. FadD can partially convert 1 mM or 2 mM OA into oleoyl-CoA (O-CoA), which can subsequently be used for synthesis of DOPE. In the absence of *fadD* template, OA can not be processed by the enzymatic pathway encoded in pGEMM7. Addition of 10 mM OA did not result in lipid synthesis, very likely because at such a high concentration OA solubilizes the LUVs. Lipid synthesis from O-CoA is less efficient in the presence of FadD which presumably can also catalyze the reverse reaction, that is the conversion of O-CoA into CoA and OA, reducing the pool of acyl donor precursor for PlsB and PlsC. Source Data are available.
